# High-accuracy SNV calling for bacterial isolates using deep learning with AccuSNV

**DOI:** 10.1101/2025.09.26.678787

**Authors:** Herui Liao, Arolyn Conwill, Ian Light-Maka, Martin Fenk, Alyssa H. Mitchell, Evan B. Qu, Paul Torrillo, Jacob S. Baker, Felix M. Key, Tami D. Lieberman

## Abstract

Accurate detection of mutations within bacterial species is critical for fundamental studies of microbial evolution, reconstructing transmission events, and identifying antimicrobial resistance mutations. While many tools have been developed to identify single nucleotide variants (SNVs) from whole-genome sequencing, they often suffer from high false positive rates due to the complexity of bacterial genomes and the need for different filtering cutoffs across sample types and sequencing depths. As datasets increase in size, the manual filtering required for high accuracy presents a significant obstacle. Here, we present AccuSNV, a novel deep learning-based tool for high-precision and automated bacterial SNV calling. Unlike traditional methods that process one sample at a time, AccuSNV leverages a convolutional neural network (CNN) that integrates alignment information across multiple samples, enhancing precision through learned across-sample patterns. We evaluated AccuSNV against seven popular SNV calling tools using simulated data from six bacterial species with varied sequencing depths, numbers of isolates, mutations, and divergence levels. To further validate its real-world utility, we tested AccuSNV on multiple curated bacterial datasets containing reported SNVs. In both simulated and real-world scenarios, AccuSNV consistently achieved the best performance. Moreover, AccuSNV provides comprehensive user-friendly downstream analysis modules and outputs, including mutation annotation information, phylogenetic inference, dN/dS calculations, and optional manual filtering. Together with the automated deep learning–based calling, these features make AccuSNV broadly accessible to users with different levels of computational expertise.

## Introduction

Single nucleotide variants (SNVs), single-base substitutions in a genome, represent the most common form of genetic variation. In bacteria, accurate identification of SNVs is a critical step in connecting genetic variation to phenotype^1,2^ and for reconstructing bacterial evolution^3^. In particular, SNV mutations form the basis of most phylogenetic inference approaches, making their accurate detection essential for epidemiology^4^. Even small errors in SNV detection can have outsized effects on downstream analyses. This is especially true for many bacterial species, which accumulate SNV mutations at a rate of only 1 - 10 SNVs per genome per year^5^. This low rate of mutation accumulation makes evolutionary inference methods sensitive to false positives, thus requiring high-precision SNV calling.

SNV calling across bacterial genomes of the same species remains a challenging task. One major source of SNV calling errors arises from alignment inaccuracies, which are exacerbated by the high diversity within bacterial species. Unlike the relatively stable and homogeneous human genome, bacteria within the same species can vary up to 5% in nucleotide identity and share only 40% of their gene content. These characteristics increase the possibility of misalignments and false variant calls^6^, particularly in low-coverage datasets or when the sample strain diverges from the reference genome^7^. In addition, cross-contamination during high-throughput sequencing preparation can lead to false positive and false negative SNV calls, posing challenges for standard mapping-based SNV filtering methods^8^. These errors can significantly reduce the precision of bacterial SNV calling.

Several SNV calling tools have been developed and applied to identify SNVs across bacterial isolates. These tools can be divided into three categories based on their underlying methodologies. The first includes probabilistic-based tools such as GATK^9^ and freeBayes^10^, which compute the likelihood of each genotype given alignment metrics such as base quality and read depth. These tools were originally designed for eukaryote organisms such as humans genomes^9^, and are not specifically optimized for bacterial genomes. The second category comprises non-probabilistic, threshold-based tools such as Samtools^11^, VarScan^12^, Breseq^13^, and Snippy^14^, which call variants by applying fixed cutoffs on parameters like read support, base quality, and strand bias. These methods are widely used due to their simplicity and informative visualizations. For example, for each predicted mutation, Breseq generates an HTML report that includes evidence pages displaying the supporting data. However, these tools require manual tuning and lack robustness across datasets with different conditions. The third category involves hybrid approaches that integrate mapping and assembly information. A representative example is BactSNP^15^, which performs *de novo* assembly of input reads and aligns the assembled contigs to the reference genome to identify SNVs. Through this approach, BactSNP aims to reduce false positives caused by read misalignment in complex genomic regions. However, the accuracy of this approach depends heavily on the quality of the assembly and the similarity between sample and reference genomes, and it remains computationally intensive for large datasets (Supplementary Table S1).

Despite their respective strengths, these existing methods share common limitations. First, they typically operate on individual isolates, which obscures across-sample patterns that can reveal alignment errors. False SNVs emerging from alignment errors exhibit distinct alignment patterns across samples (Figure 1), and failing to consider these patterns while focusing solely on single-sample evidence can lead to false-positive calls. Another common issue is reference bias: when all samples share a nucleotide that differs from the reference, methods that do not consider raw read data may erroneously call the site as polymorphic (e.g. where the reference nucleotide stems from low-coverage assembly errors), resulting in false positives. Lastly, these tools rely on manually defined thresholds or assumptions that may not generalize well, and often require extensive manual filtering to reduce false positives.

**Figure 1.**
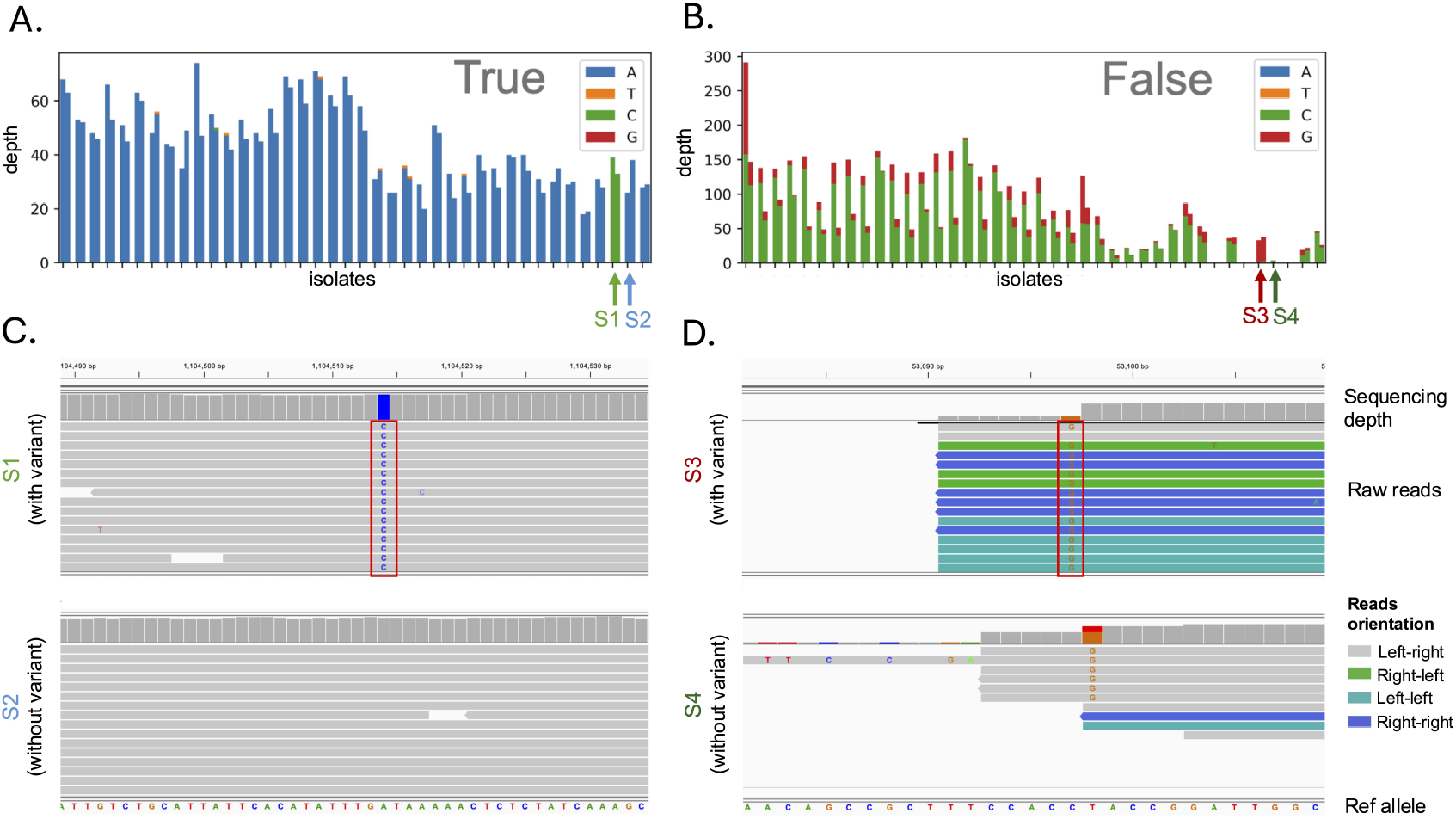
False SNVs arising from alignment errors are detectable from across-sample patterns. (A-B): Bar charts show sequencing depth across isolates for an example true SNV (A) and a false SNV (B) from real-world clinical data, with each pair of bars representing reads aligned to the forward and reverse strands for a single isolate, colored by nucleotide supported. Arrows indicate two randomly selected isolates (S1, S2 for the true SNV; S3, S4 for the false SNV) that are shown in (C) and (D). (C-D): Integrative Genomics Viewer (IGV) views of the selected representative isolates S1, S2 (C) and S3, S4 (D) carrying either the major or alternative allele for the true and false SNV, respectively. The candidate SNV is highlighted in red box. Reads are colored by pair orientation. Only left–right orientation (with the upstream read on the forward strand and downstream read on the reverse strand) is considered normal; abnormal orientations often indicate alignment errors or structural variants.

These constraints limit precision and scalability, particularly in studies involving low-depth sequencing, high strain diversity, or large-scale comparative analyses. Recent advances in deep learning^16–18^ have opened new possibilities for addressing these limitations. Unlike traditional rule-based or statistical approaches, deep learning models can automatically learn complex patterns from raw input data^19^. In the context of bacterial SNV calling, this allows models to move beyond fixed thresholds and handcrafted features, instead identifying informative signals directly from sequencing alignments. Importantly, deep learning architectures, such as convolutional neural networks (CNNs), are well-suited to capture both local alignment features and broader patterns across samples, offering a powerful framework for distinguishing true variants from false positives in diverse and noisy datasets.

In this work, we have developed AccuSNV, a deep learningbased SNV calling tool optimized for bacterial whole-genome sequencing (WGS) data. AccuSNV leverages a unique data structure that summarizes across-sample alignment information at each candidate SNV and a CNN model that derives informative patterns from this multisample read alignment data. By capturing both within-sample signals and across-sample patterns, it offers improved robustness to low-depth coverage, reference and genome divergence, and diverse realworld datasets. AccuSNV takes a reference genome and raw WGS data from multiple (*>*= 3) isolates as input, and outputs high-confidence SNVs alongside predicted probabilities, variant annotations, and informative visualizations. Through comprehensive benchmarking across both simulated and realworld datasets, we show that AccuSNV achieves consistently superior precision and generalization across a wide range of bacterial species and sequencing conditions.

## Results

### AccuSNV employs a CNN model trained on realworld bacterial isolate data

In this study, we curated real-world bacterial isolate WGS data (short reads) with manually labeled SNVs from prior studies conducted by our lab^20–22^ to serve as training and validation datasets of AccuSNV. We chose this data because: (1). real-world data contains authentic sequencing noise, technical artifacts, and biological complexity that are difficult to simulate, providing a more robust foundation for model training compared to simulated data; (2). validated SNV datasets are rare, and most publicly available ones report only true positives without false SNVs, making it impossible for models to learn the features of false calls. Our lab’s datasets overcome this limitation through manual labeling of both true and false SNVs, with standardized formats and stringent filtering (Supplementary Table S2) ensuring reliability and ease of use; (3). for these datasets, pre-extracted alignment-derived features were already available, which greatly reduced the time and computational cost of model training since raw sequencing data did not need to be reprocessed. In addition, these curated features supported the quick generation of bar charts (Figure 1) that visually distinguish high-quality from low-quality SNVs, providing an additional manual validation beyond automated filters (see Methods) and thereby enhancing label reliability; (4). these datasets encompass diverse bacterial species and experimental contexts, offering a strong founda-tion for training a generalizable model. Accession numbers and additional details regarding these WGS data are provided in Supplementary Table S3. Given these datasets, each candidate SNV site was encoded as a feature vector summarizing read-level signals and cross-sample alignment patterns, and a CNN was trained to classify them as true or false variants (Figure 2). This design leverages across-sample information to capture local sequence features and alignment patterns, forming the basis of AccuSNV’s high-accuracy SNV calling framework.

**Figure 2.**
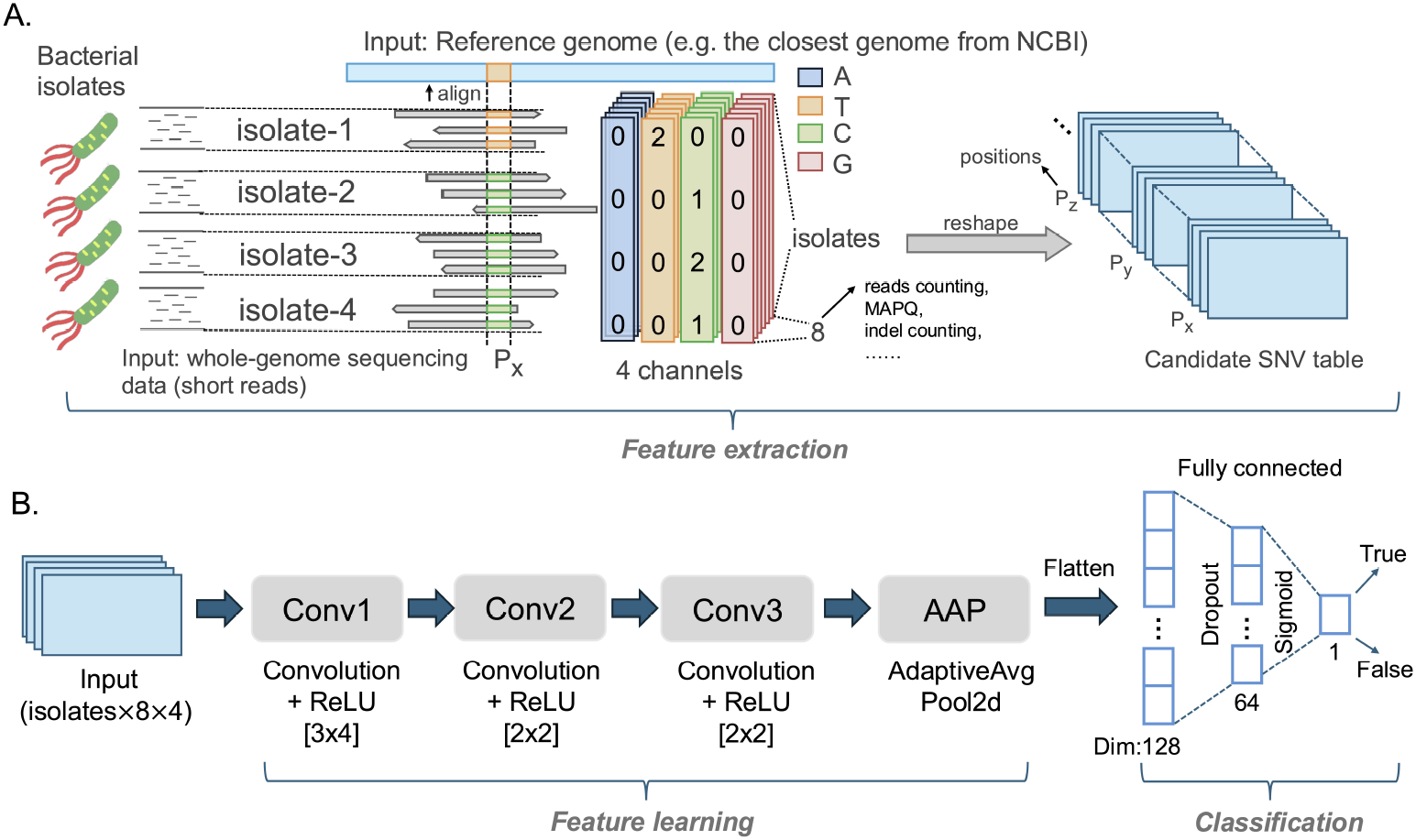
A deep learning framework that leverages across-sample comparisons for bacterial SNV calling. (A). Feature extraction pipeline: short reads from bacterial isolates are aligned to a reference genome, features at candidate SNV positions are extracted across four channels (A, T, C, G). For each channel, there are 8 features, including number of reads supporting that allele at that position on the forward and reverse reads, base qualities, and tail distances (see full list in Supplementary Figure S1). These features are reshaped into a 4D (positions × isolates × features × channels) tensor and stored in a candidate SNV table for neural network input. (B). Model architecture: the input tensor is processed through three convolutional layers with ReLU activation and varying kernel sizes (shown in brackets), followed by adaptive average pooling. The output is then flattened and passed through fully connected layers with dropout for binary classification. The model finally outputs prediction probabilities, with SNVs classified as True if probabilities *>* 0.5, otherwise False.

### The CNN model outperforms classical machine learning methods

To build AccuSNV, we first explored several classical machine learning models using the labeled SNV dataset derived from read alignments of real bacterial sequencing data from the Lieberman Lab. Input features (Supplementary Figure S1) were constructed from read alignment patterns at candidate sites along with across-sample alignment statistics, and these were used to train XGBoost, random forest, support vector machines (SVM), and logistic regression models. We compared their performance against the CNN model with two and three convolutional layers to determine the optimal model complexity for capturing local sequence context and across-sample alignment patterns. To minimize bias, we randomly split the dataset into training and validation sets five times, retrained all models on each split, and reported average performance metrics on the validation data (Supplementary Figure S2).

The CNN model consistently achieved the highest scores across all evaluation metrics on the validation dataset (Supplementary Figure S2), outperforming all classical models (p<0.05, Wilcoxon signed-rank test). Notably, the three-layer CNN also outperformed a shallower two-layer CNN architecture, demonstrating the importance of sufficient model depth for learning complex alignment patterns.

The superior performance of CNN likely stems from fundamental architectural differences between CNN and classical machine learning approaches in handling genomic alignment data. Specifically, CNN preserves the relationship between alleles and their features across samples, while classical models require flattening to 2D vectors that lose crucial across-sample patterns. Additionally, CNNs can automatically learn complex patterns from the multidimensional alignment data and detect recurring motifs (e.g. the same minor allele across many samples, Figure 1B) regardless of their position in the tensor, whereas classical models rely on hand-crafted features and cannot capture these spatial relationships. These results motivated our choice of CNN with 3 convolutional layers as the core of the AccuSNV framework.

### Overview of benchmark experiments

To evaluate the performance of AccuSNV, we benchmarked it against seven widely used SNV calling tools (GATK v.4.5.0.0, FreeBayes v.1.3.6, Samtools v.1.20, BactSNP v.1.1.0, VarScan v.2.4.6, Breseq v.0.39.0, and Snippy v.4.6.0) using both simulated and real-world datasets. The simulated data consisted of Illumina-like reads generated from six representative bacterial species across a range of sequencing depths (10X to 50X), divergence levels, and contamination scenarios, totaling thousands of samples. All tools were run using default or recommended parameters (see Supplementary Section 1). Across these diverse conditions, AccuSNV achieved consistently high accuracy, demonstrating robustness to challenges including low coverage, reference and genome divergence, and complex sequencing and alignment artifacts.

### AccuSNV achieves high precision across sequencing depths

Sequencing depth serves as a critical factor influencing the accuracy of SNV calling, where low coverage can lead to false positives or missed true SNVs^23^. In this experiment, we simulated datasets of 10 isolates with sequencing depths of 10X, 20X, 30X, 40X, and 50X to compare the performance of different tools across varying sequencing depths. First, we selected six representative bacterial species (*C. acnes, C. diffile, E. coli, K. pneumoniae, S. aureus, S. pneumoniae*) and retrieved representative reference genomes (see Methods) from NCBI^24^. To mimic real-world scenarios, we simulated datasets of closely related organisms and a more distant reference genome. As illustrated in Supplementary (Figure S3A), we introduced 1% SNVs and 500 indels into each reference genome with SimuG^25^ to create a root genome, which serves as the common ancestral sequence from which all simulated isolates will subsequently diverge. From this, we generated 10 mutant genomes (with about 70 mutant positions introduced per genome on average) per species with a phylogenetic structure intended to mimic realistic evolutionary relationships and simulated Illumina reads at 10X–50X coverage using ART^26^, resulting in 300 samples (10 isolates × 6 species × 5 depths). All reads were aligned back to the corresponding reference genome, SNV calling was performed using each tool, and precision, recall, and average F1 scores of each tool was calculated (Figure 3).

**Figure 3.**
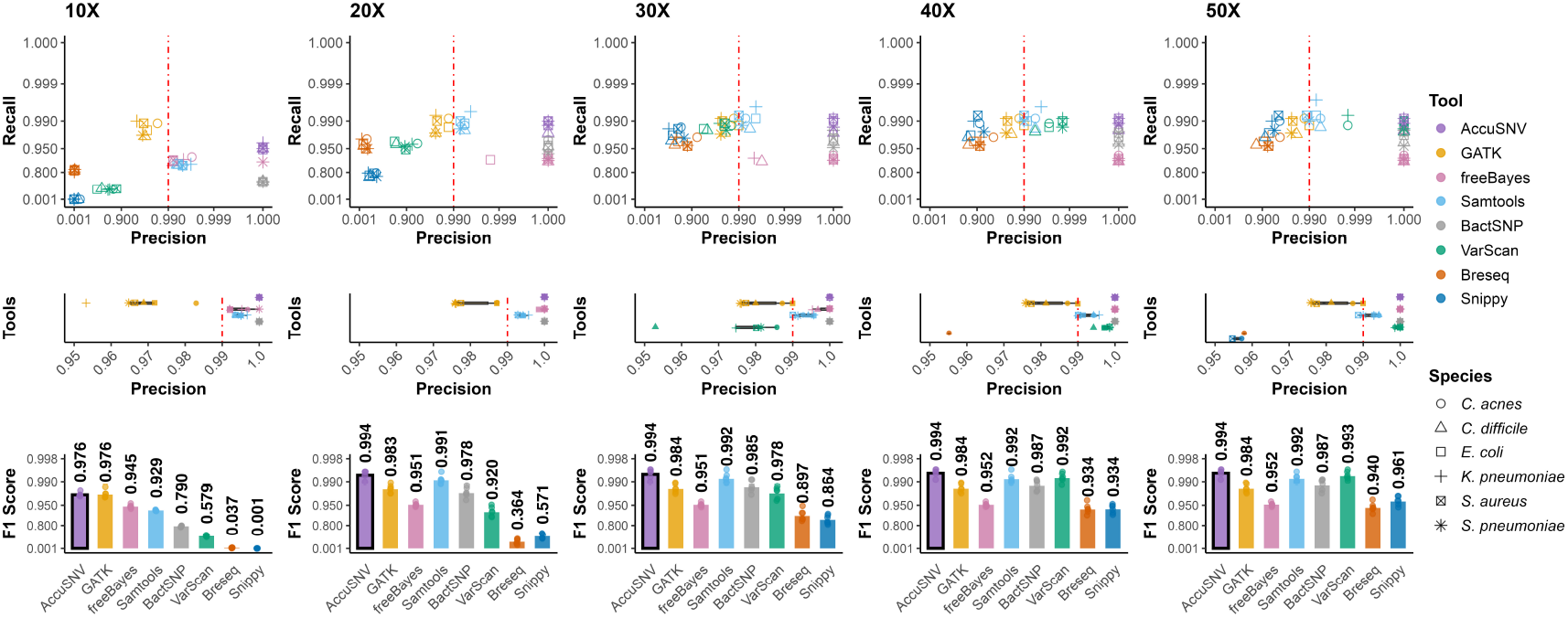
AccuSNV achieves high precision across simulated datasets with different sequencing depths. Precision, recall, and average F1 scores of AccuSNV and seven other variant-calling tools were evaluated across simulated bacterial datasets at five sequencing depths (10X, 20X, 30X, 40X, and 50X). Axes use a transformed scale (− log_10_(1.0001 − *x*)) to enhance visualization in the higher range (0.9-1). Shapes represent species and colors represent tools. The vertical line at 0.99 precision highlights how more tools achieve high performance at greater sequencing depths. Top panels show precision–recall scatter plots for all tools and species at each depth. Middle panels provide a zoomed-in view of precision distributions across tools and species. Bottom panels summarize the average F1 score for each tool. The bar corresponding to AccuSNV is highlighted by a black outline.

The performance of all tools improves with increasing sequencing depth, highlighting the importance of high-depth sequencing data for existing tools to accurately identify bacterial SNVs. In addition, AccuSNV achieves the highest average F1 score across all tested datasets (Figure 3). In particular, AccuSNV achieves F1 scores of 97.6% and 99.4% at coverages as low as 10X and 20X, respectively, while maintaining 100% precision across all experiments. Among other tools, BactSNP also achieves perfect precision but with lower recall compared to AccuSNV. At a sequencing depth of 10X, GATK has higher recall than AccuSNV, but its average precision is 96.8%, compared to 100% for AccuSNV. Samtools and VarScan perform competitively at higher depths but underperform at 10X, with F1 scores dropping to 92.9% and 57.9%, respectively. The remaining three tools, freeBayes, Breseq, and Snippy show poorer performance compared to other tested tools across all datasets. The performance of Breseq and Snippy drops markedly at 10X depth, because the conservative filters, such as minimum depth, allele-fraction and strand-balance requirements, and evidence-count thresholds, discard many true variants once the effective coverage falls below these cutoffs.

### AccuSNV maintains high accuracy on large and noisy simulated datasets

In real-world scenarios, sequencing samples from different isolates often have varying depths and complex phylogenetic relationships. Moreover, previous studies have shown that the divergence between the target genome and the reference genome significantly impacts bacterial SNV calling performance^7^. Lastly, many real-world datasets contain large isolate collections, introducing additional complexity to SNV detection. To better reflect these real-world challenges, we generated an additional large-scale, noisy simulated dataset for evaluation. Notably, freeBayes and Breseq were excluded from this experiment and subsequent real-world data evaluations due to their lower accuracy in earlier evaluations and substantially longer runtimes (Supplementary Table S1).

To investigate the influence of the reference genome, we first generated four root genomes per species of varing reference divergence from the reference genome. From each root genome, we simulated a phylogenetic tree with 100 strains using Msprime^27^, which provided the number of mutations along each branch for the given mutation rate (used 5 × 10^−10^ here). We then assigned 80% of the mutations as SNVs and 20% as indels, and used SimuG to generate the corresponding mutated genomes. We focused solely on SNVs and indels without larger recombination or rearrangement events, as this analysis targets very closely related isolates where such structural variations are rare. Paired-end reads were simulated at random depths between 15X and 70X. In total, this yielded 2,400 samples (100 isolates × 6 species × 4 reference divergence levels) for evaluating SNV calling tools (Figure S3B).

AccuSNV exhibits the most competitive performance among all tested tools, with average F1 scores exceeding 99% across all datasets (Figure 4). It maintains perfect precision across all reference divergence levels and achieves an F1 score above 99.1% even at the highest reference divergence (1%). In contrast, other tools show greater variability: VarScan, Samtools, and Snippy all experience significant drops in precision or recall at higher reference divergence levels. Although GATK performs comparably to, or slightly better than, AccuSNV at the 0.05% and 0.1% reference divergence levels in terms of F1 score, this is driven by increased recall at the cost of reduced precision, which is suboptimal when comparing closely-related genomes where precision is critical^28^. Bact-SNP also maintains perfect precision across all datasets but with consistently lower recall, leading to reduced F1 scores, especially under higher reference divergence.

**Figure 4.**
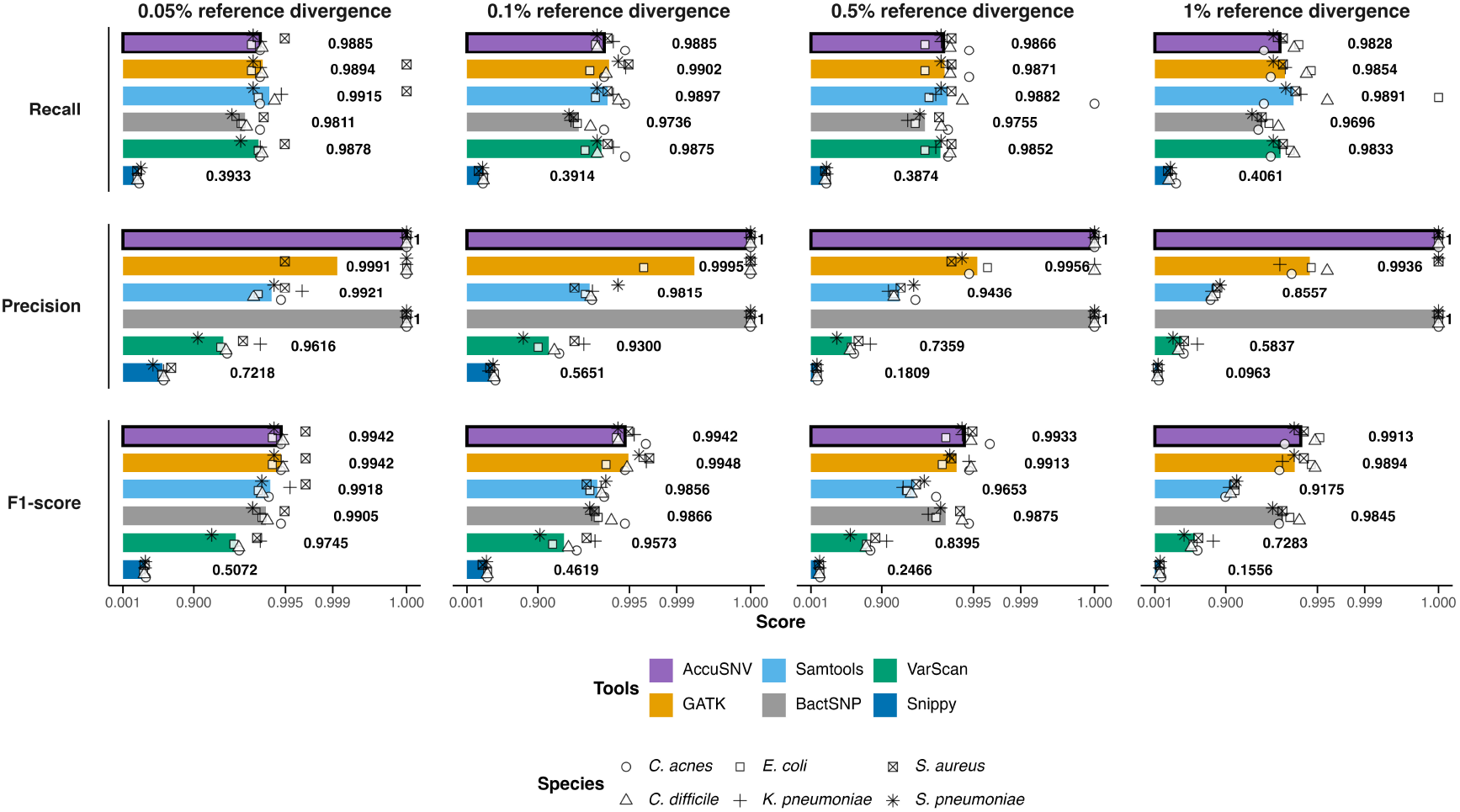
AccuSNV maintains high accuracy on large and noisy simulated datasets. Performance comparison of six SNV calling tools across four reference divergence levels. Axes use a transformed scale (−log_10_(1.0001 − *x*)) to enhance visualization in the higher range (0.9-1). Shapes represent species and colors represent tools. Shapes represent species and colors represent tools. Panels show average recall (top), precision (middle), and F1 score (bottom) across all species. AccuSNV bars are highlighted with a black outline.

Overall, all tools exhibit decreased performance as the divergence between the root genome and the reference genome increases, highlighting the growing difficulty of variant calling when read-reference mismatches become more prevalent. Despite this, AccuSNV shows minimal performance fluctuation across reference divergence levels. In addition, tools such as Samtools and VarScan, which performed well in the previous experiments with only 10 strains at 50X, uniform sequencing depth, and simplified phylogenetic structure, showed markedly reduced performance on this more realistic and noisy dataset. These results indicate that the robustness of existing tools may decline under more complex and realistic sequencing conditions. In contrast, AccuSNV maintains both high accuracy and consistency, supporting its use in largescale bacterial variant detection across diverse and complex conditions.

### Experiments on highly variable isolates

Some real-world bacterial populations, such as those from environmental samples, can exhibit substantial genetic variability between isolates. To evaluate how well different tools perform under such conditions, we varied the mutation rate in 100 *E. coli* genomes using Msprime (see Supplementary Figure S3B), based on a root genome containing 1% sequence variation. Specifically, we tested mutation rates of 5 × 10^−9^, 5 × 10^−8^, and 1 × 10^−7^, along with the original rate of 5 × 10^−10^ per site per generation. We reran all tools on these datasets and recorded the results (Supplementary Figure S4).

As mutation rates increased, all tools exhibited more false positives and false negatives. Nevertheless, AccuSNV achieved a high average F1 score (98.85%) by balancing recall and precision. GATK (with 98.81% average F1 score) showed the most competitive F1 score at higher mutation rates but had lower precision than AccuSNV. In contrast, BactSNP (with 98.11% average F1 score) maintained higher precision across all datasets, but its recall was substantially lower. Samtools (with 98.33% average F1 score) achieved the best recall across all datasets, yet it produced more false positives at higher mutation rates. Overall, AccuSNV demonstrated the highly robust and reliable performance across mutation rates, underscoring its effectiveness for analyzing highly variable bacterial isolates.

### Computational efficiency comparison

To compare the running time and memory usage of different tools, we applied all tools to 10-strain and 100-strain *E. coli* simulated datasets and recorded the computational performance metrics (Supplementary Table S1). AccuSNV demonstrated competitive efficiency, completing analysis in 30.3 minutes for 10 isolates (50X) and 64.5 minutes for 100 isolates (15X-70X), with memory usage of 1,140 MB and 1,340 MB respectively. While some tools like Snippy showed faster execution times (5 minutes on a 10-strain dataset and 50 minutes on a 100-strain dataset), AccuSNV maintained better performance without the extreme computational demands observed in tools like freeBayes and Breseq, which failed to complete within reasonable time limits on the larger dataset (running more than 2 days on the 100-strain dataset). The results demonstrate that AccuSNV provides a balance between accuracy and computational efficiency for bacterial SNV calling applications.

### AccuSNV shows good robustness in simulated contaminated samples

To assess the robustness of SNV calling tools under contamination, we simulated datasets with increasing levels of contamination, either from other strains of the same species or from closely related species, under both high and low sequencing depths. AccuSNV consistently achieved higher F1 scores than other tools in low-depth contaminated datasets and maintained perfect precision in all tested datasets. In high-depth settings (50X depth) with contamination from both other strains and closely related species, VarScan achieved the highest F1 scores across all datasets, with AccuSNV ranking second due to slightly lower recall. However, in this high-depth setting, when contamination originated from closely related species, VarScan exhibited reduced precision compared to AccuSNV. Full results are provided in Supplementary Section 2.1, Figure S6-S9.

### AccuSNV demonstrates high accuracy on real-world test data

Real-world data from independent studies, featuring diverse sequencing protocols, depths, species, and collection contexts, offer a more rigorous assessment for evaluating the robustness and accuracy in practical applications, as simulations rarely include all types of error and variability in the real world. In this experiment, we evaluated AccuSNV and other tools using four publicly available bacterial sequencing datasets^29–32^, each derived from a distinct study, author group, and accompanied by manually curated and reported SNVs that underwent careful quality control rather than being taken directly from automated tool outputs. The four datasets differ in read types (single-end vs. paired-end), bacterial species, and isolate numbers (Figure 5). The dataset from Snitkin et al. (2012) comprises single-end reads from a hospital outbreak of *Klebsiella pneumoniae*, while the others involve paired-end sequencing of *Staphylococcus aureus* or *Clostridioides difficile* collected under varying evolutionary or clinical contexts. These differences in sequencing protocols, strain diversity, and sampling settings create a heterogeneous and realistic benchmark for evaluating SNV-calling accuracy and robustness.

**Figure 5.**
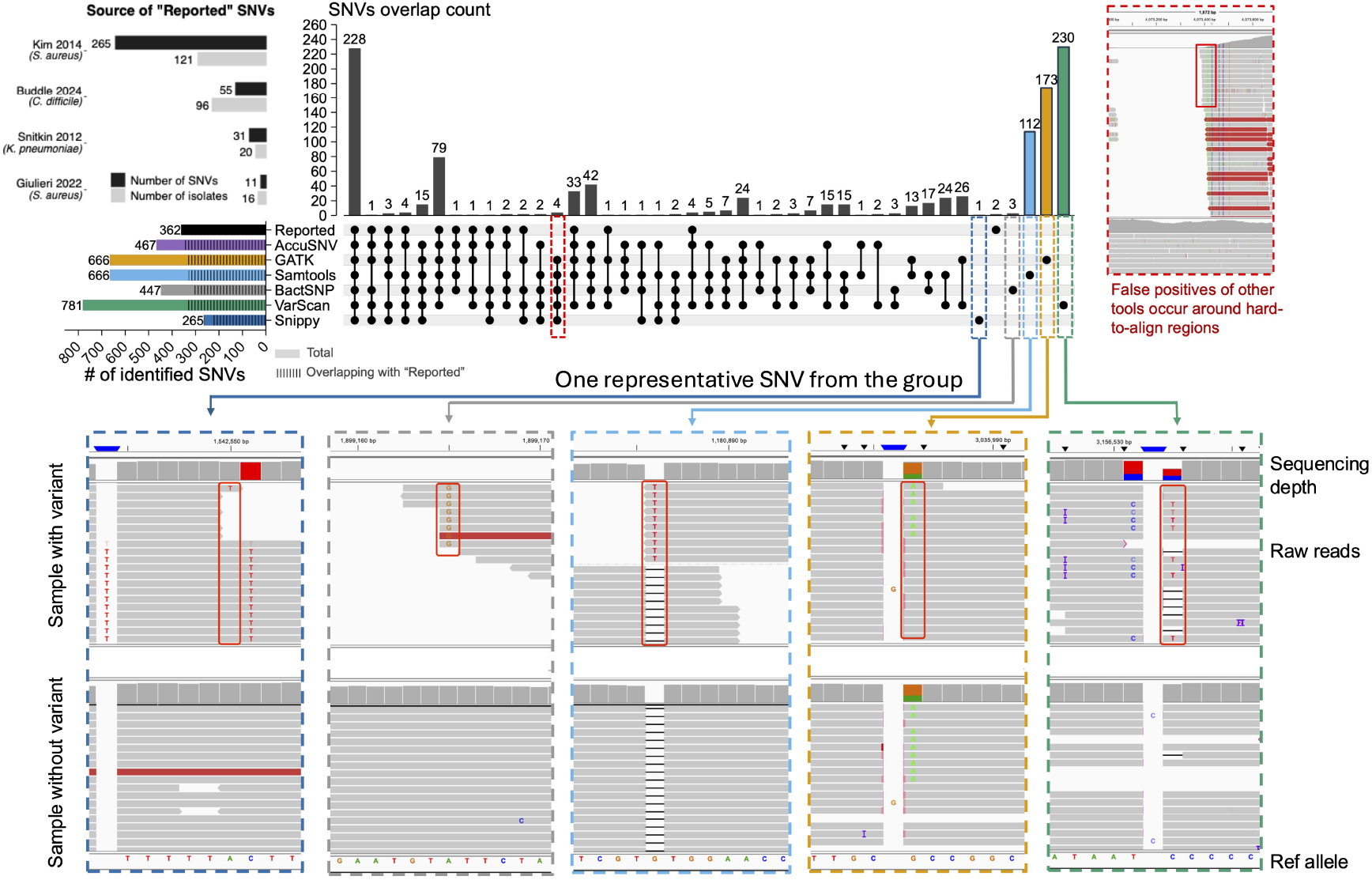
AccuSNV demonstrates high accuracy and precision on additional real-world bacterial sequencing datasets. We evaluated SNV concordance between AccuSNV and five other tools across four published bacterial datasets with curated SNVs (top left panel). UpSet plots (center) display the SNVs overlap count of tool-identified SNVs and reported SNVs. The red-outlined dash box highlights a group of four SNVs identified by all tools except AccuSNV and the original study (corresponding IGV^33^ screenshot is shown in the top right panel). The IGV screenshot includes two sample tracks, one carrying the reference allele and one carrying the alternative allele, with the variant position highlighted by a red box. Red reads indicate an inferred insert size larger than expected, which may suggest the presence of a deletion or a possible alignment artifact. The bottom panels display IGV screenshots of five unique SNVs identified by other methods but not supported by AccuSNV or the original study. The colored borders around each IGV screenshot correspond to the dashed boxes in the UpSet plot, indicating the group from which each SNV originated. These screenshots highlight typical error signatures such as misalignments near read starts/ends, strand bias, soft-clipped bases, and low-complexity sequences, etc (as reported in Koboldt et al (2020)^34^).

All tools called distinct sets of SNVs across these data sets, with none matching the reported SNVs exactly (Figure 5). Across all four datasets, AccuSNV demonstrates strong concordance with reported SNVs while minimizing tool-specific SNVs unsupported by other methods or the original studies (only 110 unreported SNVs, compared to 127-422 for other tools, except Snippy, which had 26 but identified much fewer reported SNVs).

Inspection of discordances among SNV calling methods illustrates that AccuSNV avoids false positives called by other tools, generally caused by alignment errors. For example, a noteable group of SNVs is highlighted in the red dashed box (Figure 5); these four SNVs from the same study were called by all other tools but not called by AccuSNV or the original study. To investigate the basis of this discrepancy, we visualized the corresponding BAM files of these four SNVs, one of which is shown in the panel outlined on the right. These four SNVs appear to be false positives located in hardto-align regions, due to divergent regions between the sample and reference.

In addition, all tools except AccuSNV reported unique SNVs that were not supported by any other method or the original study. To further examine these unique SNVs, we manually reviewed five random unique SNVs from tested tools and visualized their read alignments (bottom panel of Figure 5). In all cases, the alignments revealed problematic mapping features, such as soft clipping, inconsistent base support, or alignment gaps. These findings suggest that these unique SNVs called by other tools are false positives resulting from alignment errors. This observation agrees with previous studies, such as the one by Koboldt (2020)^34^, which identified similar artifacts in short read alignments leading to false-positive variant calls. AccuSNV is able to filter these false positives by capturing informative across-sample features from alignment data, rather than relying on manually defined thresholds or manual filtering of individual positions.

### Output and downstream analysis modules of AccuSNV

To support users with varying levels of computational expertise, AccuSNV offers comprehensive output and downstream analysis modules based on the identified SNVs. The core output includes a compressed SNV table in NumPy’s `.npz` format (a compact binary file for storing arrays), a human-readable tab-separated summary of all the SNVs (TSV), a graphical HTML report, and a set of quality control (QC) figures (Figure 6).

**Figure 6.**
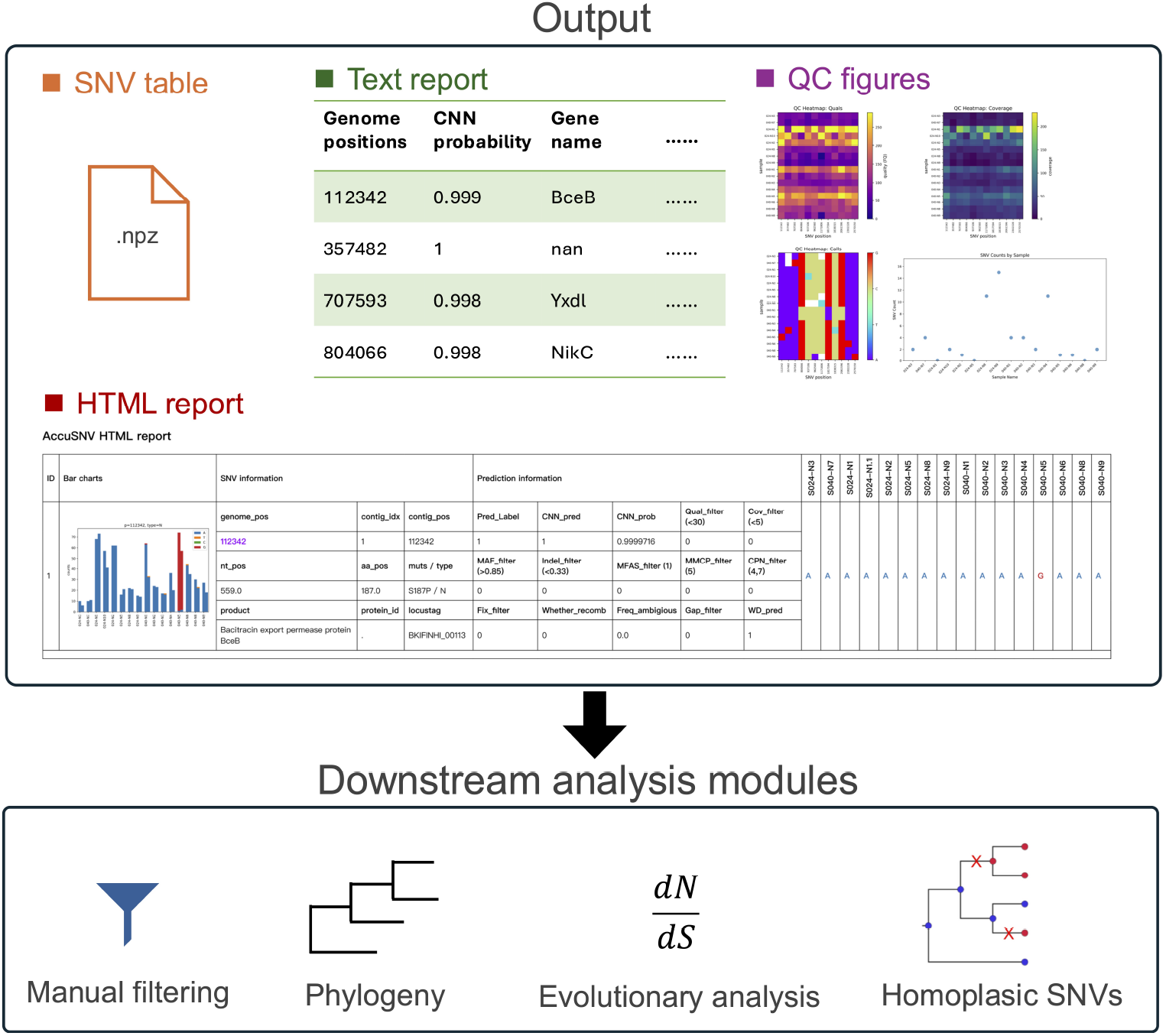
Output and downstream analysis modules of AccuSNV. The core outputs include: (1). a compressed SNV table in NumPy’s `.npz` format storing detailed feature vectors and prediction scores, (2). a human-readable text summary file (TSV) listing all identified SNVs with key attributes, (3). a set of quality control (QC) figures summarizing read coverage, base calls, mapping quality scores, etc, (4). a HTML report that integrates summary tables and bar charts of identified variants. Homoplasic SNVs in the figure refers to variants where the same derived nucleotide arises independently in two or more lineages since their divergence from a common ancestor with a different ancestral base. These SNVs can be used to infer parallel, convergent, or revertant evolutionary events^35^.

The SNV table stores all variant-level features and prediction outputs in an efficient binary format, which serves as input for downstream modules. The text summary presents essential information for each SNV, including the genomic position, predicted probability, and annotation information, etc. This enables rapid inspection and integration with external pipelines. The HTML report provides an interactive platform for variant review. Each predicted SNV is visualized with a bar chart showing allele frequency distributions across isolates (as in Figure 1), alongside detailed metadata such as strand-specific depth, mapping quality statistics, and various filter flags. A demo output HTML report can be found at https://heruiliao.github.io/. This design helps users assess variant confidence and identify potentially ambiguous cases that may require manual curation. In addition, AccuSNV generates multiple QC figures to assist in data quality assessment, including heatmaps of genome coverage and allele count distributions across samples.

To facilitate related applications, AccuSNV includes builtin modules for common downstream analyses. These modules enable manual filtering, phylogenetic tree construction, calculation of *d*_*N*_*/d*_*S*_ ratios to assess selective pressures, and identification of homoplasic SNVs^35^. Additionally, users can export filtered variants for phylogenetic analysis with external tools.

Together, these outputs and utilities make AccuSNV a versatile and accessible tool for high-confidence bacterial SNV detection and interpretation.

## Discussion

In this study, we present AccuSNV, a deep learning-based variant calling framework designed for high-precision single nucleotide variant (SNV) detection from bacterial wholegenome sequencing (WGS) data. By encoding multisample read alignment data into structured feature vectors and leveraging a convolutional neural network (CNN), AccuSNV effectively captures across-sample patterns that are often ignored by conventional methods. Our approach addresses key challenges in bacterial variant calling, including low sequencing depth, high intraspecies diversity, and genome complexity. As a result, AccuSNV enables accurate bacterial SNV calling for users with varying levels of expertise, without the need for hard-coded thresholds or extensive manual filtering. While a few tools occasionally achieved comparable or better performance under specific conditions, AccuSNV was consistently the best or among the top-performing methods across all sequencing depths, reference and sequence divergence levels, and contamination scenarios, demonstrating its broad applicability and scalability for bacterial genomics research.

Despite these advances, AccuSNV has several limitations. First, misclassfications may still arise in extremely challenging or ambiguous cases encountered in real-world data, especially those involving false-positive SNVs arising from an error mode not represented in the training set. To mitigate this, AccuSNV provides an interactive HTML report and multiple quality control visualizations that allow users to inspect candidate SNVs and flag potential false positives for downstream curation. Second, AccuSNV is specifically designed for SNV calling and does not currently support the identification of structural variants (SVs), such as insertions, deletions, or inversions. Third, the method is not directly applicable to metagenomic datasets, where a single sample may contain multiple closely related strains, resulting in more complex and ambiguous alignment patterns than those encountered in isolate-based WGS.

We note that in all of our evaluation experiments, we assess each genomic position as either a true or false variant site, rather than evaluating individual per-sample variant ep. Thus, many individual false calls are reduced to only a single count in our evaluations. In practical applications, especially under low coverage or high reference divergence many samples may have false-positives or false-negatives at a single position. As such, the position-level evaluation presented here provides a conservative benchmark of tool performance in these noisy and complex scenarios.

Looking forward, several directions can extend the utility of AccuSNV. One promising avenue is to expand the framework to support small indel detection by incorporating indel-aware features and labels during model training. Another direction is to explore whether across-sample alignment patterns in metagenomic data can be modeled to distinguish strain-specific SNVs, potentially enabling AccuSNV to oper-ate in complex microbial communities. However, these efforts may be constrained by the limited availability of real-world datasets with manually validated labels.

In conclusion, by reducing the need for manual thresholding and incorporating interpretable quality controls, AccuSNV makes high-confidence bacterial SNV calling accessible to users with varying levels of computational expertise. This accessibility makes it particularly well suited for studies of within-microbiome evolution, large-scale evolution experiments, and bacterial epidemiology, even when working with low-coverage or noisy sequencing data. By lowering technical barriers, AccuSNV enables broader adoption of high-precision SNV calling in diverse research and clinical settings.

## Methods

### Overview of AccuSNV

AccuSNV is designed for high-precision single nucleotide variant (SNV) calling from whole-genome sequencing data of bacterial isolates. It takes a reference genome and wholegenome sequencing short-read data from multiple bacterial samples (isolates) as input, and outputs identified SNVs along with associated information such as annotations, predicted probabilities, and visualization results, etc (Figure 2 and 6). AccuSNV leverages deep learning to distinguish true variants from sequencing or mapping artifacts, eliminating the need for manual filtering. Specifically, to address challenges such as low sequencing depth, genomic intraspecies diversity, variability in sample size, and sequencing quality across datasets, AccuSNV employs a CNN-based framework to learn informative patterns from multi-sample read alignment data. Here, we chose CNN for its ability to extract local features and maintain translation invariance^36^, which enables the robust detection of across-sample patterns from diverse and noisy bacterial isolate alignments. To implement this CNN-based approach, AccuSNV first identifies candidate SNV sites using Samtools and extracts read-level features from bacterial isolate alignments using a Snakemake-based pipeline (Figure 2A), including reads orientation, reads counting, mapping quality score, etc (Supplemetary Figure S1). These features are converted into a four-dimensional numerical feature vector, allowing the CNN to better capture both local and across-sample patterns. The CNN model (Figure 2B) was trained using these four-dimensional feature vectors as input, and the final pretrained model is provided to perform binary SNV classification on new datasets.

### Reference genomes used in this study

*Cutibacterium acnes* (*C. acnes C1*, GCF_000302515.1), *Clostridium difficile* (*C. diffcile R20291*, GCF_000027105.1), *Escherichia coli* (*E. coli str. K-12 substr. MG1655*, GCF_000005845.2), *Klebsiella pneumoniae* (*K. pneumoniae subsp. pneumoniae NTUH-K2044*, GCF_000009885.1), *Staphylococcus aureus* (*S. aureus subsp. aureus NCTC 8325*, GCF_000013425.1), *Streptococcus pneumoniae* (*S. pneumo-niae R6*, GCF_000007045.1) genomes were selected as reference genomes in this study.

### Training and validation datasets

To train and evaluate the model, we collected a total of 7,638 “quality-filtered” SNVs (labeled as “True”) from 5,432 isolates from three datasets published in previous studies^20–22^ conducted by our lab, and 6,306 “low-quality” SNVs filtered out during processing for these studies (labeled as “False”). These datasets comprise 222 lineages (finely resolved clades separated by fewer than 100 mutations across the core genome^21,22^) and include carefully curated SNVs based on extensive manual validation of filters in a study-specific manner. The criteria used in each study to define “quality-filtered” SNVs are summarized in Supplementary Table S2. To verify the collected positions in the training data, we randomly picked up to 10 true and 10 false positions per lineage (for a maximum of 20 per lineage), but some lineages contained fewer than 20 available sites, resulting in 2,859 positions that were manually inspected using generated bar charts (see example in Figure 1). These bar charts are available at https://zenodo.org/records/17058222. Manual inspection confirmed that the assigned labels were accurate and consistent with the expected patterns. These data were split into training and validation sets at an 4:1 ratio, strat-ified by lineages across the three datasets. This ensured that the validation set contained isolates from different lineages and datasets, resulting in 10,887 SNVs for training and 3,057 SNVs for validation, and thereby improving the generalizability of the trained model. Accession numbers and additional details regarding the training data are provided in Supplementary Table S3. Additionally, to assess tool performance on real-world data from independent studies, we collected 362 reported SNVs from four different published datasets, where SNVs had been carefully validated in the original studies through stringent filtering and manual inspection rather than taken directly from automated tool outputs. Detailed information on all seven datasets is provided in Table 1.

**Table 1.** Summary of the seven real bacterial whole-genome sequencing datasets used in this study. The first three datasets were used for model training and validation, while the remaining four served as independent test sets for additional evaluation. SNV calling was performed within lineages, defined as closely related clades identified based on core-genome similarity^21,22^. “-” indicates no reported false SNVs in the original study.

| Datasets ID | Type | Species | # of lineages | # of isolates | # of True SNVs | # of False SNVs |
| --- | --- | --- | --- | --- | --- | --- |
| Zhao_2019<sup>20</sup> | Training/Validation | *B. fragilis* | 11 | 556 | 227 | 492 |
| Conwill_2022<sup>21</sup> |  | *C. acnes* | 50 | 854 | 1,972 | 3,840 |
| Baker_2025<sup>22</sup> |  | *C. acnes* | 85 | 2,007 | 3,444 | 554 |
|  |  | *S. epidermidis* | 76 | 2,015 | 1,995 | 1,420 |
| Kim_2014<sup>31</sup> | Test | *S. aureus* | 1 | 121 | 265 | - |
| Buddle_2024<sup>32</sup> |  | *C. difficile* | 1 | 96 | 55 | - |
| Snitkin_2012<sup>29</sup> |  | *K. pneumoniae* | 1 | 20 | 31 | - |
| Giulieri_2022<sup>30</sup> |  | *S. aureus* | 1 | 16 | 11 | - |

### Identification of candidate SNVs

To identify candidate SNV sites from raw sequencing data, we implement a modular pipeline built with Snakemake^37^. The process begins with quality trimming of reads using Cutadapt (v.1.18)^38^ and Sickle^39^ (v.1.33; -g -q 20 -l 50 -x -n), followed by reference-based alignment via BWA-MEM^40^ (v.0.7.18). Aligned reads are converted to sorted and indexed BAM files. SAMtools markdup (v.1.21.1; -r -s -d 100 -m s) is used to remove duplicate reads from each sample. Pileup and VCF files are then generated using Samtools^11^ mpileup (v.1.21.1; -q30-x -s -O -d3000), and Bcftools^41^ view (v.1.21.1; -Oz -v snps -q.75). Enabled by the parallel management of independent tasks via Snakemake, these computationally intensive steps can be completed efficiently, even across large-scale isolate collections (Supplementary Table S1).

For each input sample, preliminary SNVs are extracted from VCF files and saved in compressed intermediate files. Then, the pipeline aggregates variant positions across all samples, and combines these positions with alignment-derived metrics, such as read counting and mapping quality score, into a candidate mutation table using custom python scripts. This table contains informative alignment statistics for each candidate SNV and serves as the input for downstream analysis.

### Feature extraction

To ensure CNN can effectively capture informative patterns across bacterial isolates, AccuSNV transforms raw alignment data into structured four-dimensional feature vectors. Specifically, for each candidate SNV position, we aggregate information from all reads supporting each allele in each sample, including base counts on forward and reverse strands, mapping quality scores, and indel signals (the number of reads supporting insertions and deletions, broadcast across the four base channels per sample; Supplementary Figure S1). Notably, these features were selected because they are commonly used in the manual filtering step across previous studies^5,20,21^ for distinguishing “quality-filtered” SNVs from “low-quality” positions. In addition, we incorporate two normalized coveragebased features, 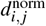 and *r*_*i, j*_, to better represent both across-position and across-sample variation. These two features are computed separately for forward and reverse strands to retain strand-specific information:

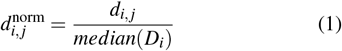

where *d*_*i, j*_ is the raw read depth of sample *i* at site *j*, and *median*(*D*_*i*_) is the median depth across all positions in sample *i*.

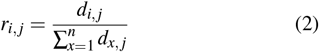

Here, *r*_*i, j*_ refers to the proportion of read depth from sample *i* relative to the total depth across all *n* samples at position *j*, serving as a relative coverage signal across isolates.

During feature extraction, we further observed that certain low-quality positions display extremely unbalanced coverage distributions, where the depth of the putative minor allele is orders of magnitude lower than that of the major allele (see Supplementary Figure S5). Such unusual “gap cases” are underrepresented in the training data and tend to be misclassified by the CNN. To ensure these extreme cases are consistently represented, we introduced a normalization-based adjustment in the feature encoding step, implemented through a z-score framework:

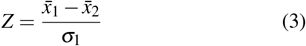

where 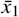 and 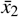 denote the mean normalized read depths (*di, j*^norm^) of the major-allele–supporting and alternateallele–supporting samples, respectively, and *σ*_1_ is the standard deviation of normalized read depths among the majorallele–supporting samples. The one-sided p-value is then calculated as:

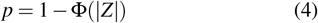

where Φ denotes the cumulative distribution function of the standard normal distribution. Positions with *p <* 0.01 are flagged as gap cases, and the corresponding feature values are normalized to ensure consistent representation across isolates. This adjustment is fully integrated into the feature construction process, making the AccuSNV more robust to highly skewed depth profiles. We also conducted an ablation study on this normalization-based adjustment to evaluate it’s impact on performance (Supplementary Section 2.2). The results show that this design improves the robustness of AccuSNV on datasets containing positions with extreme coverage imbalance by reducing false positives while maintaining high recall (Supplementary Figure S4).

These features are then encoded as a four-dimensional feature vector (Figure 2), where the last dimension represents the nucleotide channels. This four-dimensional feature encoding has two key advantages over traditional feature representation like raw VCF statistics or flattened alignment matrices. First, it encodes comprehensive alignment features in a structured, image-like format that is well suited for the CNN model, enabling it to leverage translation invariance and local feature extraction. Second, it preserves across-sample information, enabling the model to capture shared or contrasting patterns among isolates at each site. These patterns are not accessible to conventional single-sample variant callers but are essential for distinguishing true SNVs from sequencing or alignment artifacts.

### Deep learning framework

Given the extracted feature vectors, we employ a convolutional neural network to classify the candidate SNVs. There are two major challenges in applying CNN to this task. First, the number of bacterial isolates varies across datasets, leading to input tensors with different dimensions along the sample axis. Such variability poses a problem for standard CNN architectures, which require fixed input dimensions for training and inference. Second, compared to natural image tensors that often contain rich spatial information and high-dimensional features, our input tensors have a much lower feature dimensionality. Commonly used deep or parameter-heavy models are therefore prone to overfitting, especially given the limited number of labeled training examples.

To address these issues, we designed a CNN architecture to accommodate the varying alignment data. The model consists of convolutional, pooling, and fully connected layers (Figure 2B). Specifically, we used three convolutional layers to hierarchically capture both localized signal structures and broader across-sample patterns. These three layers use 32, 64, and 128 filters with kernel sizes of 3 × 4, 2 × 1, and 1 × 1, respectively. Each convolutional layer is followed by a ReLU activation function. To accommodate inputs with variable sample sizes, we apply an adaptive average pooling operation^42^ along the sample axis, compressing variable-length inputs into a fixedsize representation. Unlike standard pooling layers with fixed kernel sizes, adaptive pooling dynamically divides the input into regions such that the output always has the same predefined shape, regardless of the input size. Let the **X** ∈ ℝ ^*x×y×z×*4^ be the input feature vector, where *x* is the number of candidate SNV postions, *y* is the number of samples (bacterial isolates), *z* is the number of features, and 4 is the input channels. Then, the output tensor **Z** of the adaptive average pooling layer is:

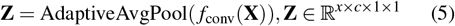

where *c* is the number of output channels from the last convolutional layer. This operation enables the model to generalize across datasets with different sample sizes while learning informative patterns across isolates. The output of the pooling layer is fed into a fully connected layer with 64 units, followed by dropout (*p* = 0.5) and a final sigmoid output node that produces the probability of the SNV being true. By default, the classification cutoff is 0.5.

### Model training

The model was trained using real-world bacterial wholegenome sequencing data with curated labels obtained from previously published studies^20–22^, as summarized in Table1. During training, the binary cross-entropy loss function was used as the objective function.

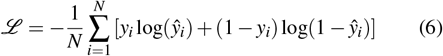

where *y*_*i*_ ∈ {0, 1} is the true label of the *i*-th candidate SNV and *ŷ*_*i*_ ∈ (0, 1) is the predicted probability. We implemented the model using PyTorch, and trained it using the Adam optimizer with a learning rate of 0.0001, a batch size of 32, and early stopping based on validation loss. The model was trained for 150 epochs, and the one with the lowest validation loss was saved for all classification tasks in our evaluation experiments. The learning curve of the model training is shown in Supplementary Figure S10.

### Evaluation metrics

To evaluate the performance of different tools, we used five standard metrics: accuracy, precision, recall, F1 score, and area under the curve (AUC). Let *TP, FP, TN* and *FN* denote the number of true positives, false positives, true negatives, and false negatives, respectively. The evaluation metrics are defined as:

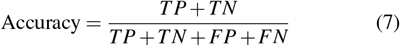

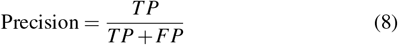

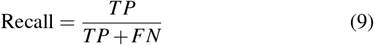

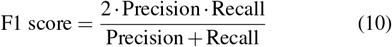

AUC was calculated based on different models’ predicted probabilities and reflects their ability to distinguish true SNVs from false ones across different decision thresholds.

## Code and data availability

The source code of AccuSNV is freely available at https://github.com/liaoherui/AccuSNV. AccuSNV is also available via Bioconda at https://anaconda.org/bioconda/accusnv. Datasets used in this paper are all publicly available. Bar charts for the 2,859 positions selected for manual label inspection are available at https://zenodo.org/records/17058222. A demo output HTML report of AccuSNV can be found at https://heruiliao.github.io/.

## Acknowledgements

This work has been supported by MIT-Novo Nordisk Artificial Intelligence Postdoctoral Fellowship to HL, NIH grant 1DP2GM140922 and R35GM156282 to TDL. ILM, MF, and FMK were supported by the Max Planck Society. During the preparation of this work, the authors used ChatGPT (https://chatgpt.com/) in order to rephrase sentences. We thank members of the Lieberman Lab and Key Lab for helpful comments.

## Author contributions statement

Methodology: HL, PT, AC, EQ, AHM, JSB, MF, ILM, FMK, TDL; Investigation: HL, AHM, TDL; Funding acquisition: HL, TDL; Writing: HL, TDL.

