## Supplementary file 1 for "High-accuracy SNV calling for bacterial isolates using deep learning with AccuSNV"

September 30, 2025

#### 1 Supplementary Methods

In this section, we provided the command lines for running all bacterial SNV callers in the main article.

##### 1.1 AccuSNV

```
python accusnv_snakemake.py -i <input_csv> -r <ref> -o <out>
```

##### 1.2 GATK

```
gatk HaplotypeCaller -R <ref> --emit-ref-confidence GVCF -I <input.bam> -O <out.gvcf>

gatk CombineGVCFs -R <ref>  --variant <out1.gvcf> <out2.gvcf> ... -O <merged.vcf>

gatk --java-options "-Xmx4g" GenotypeGVCFs -R <ref> -V <merged.vcf> -O <merged.gvcf>

gatk VariantFiltration --filter-expression "QD<2.0||MQ<40.0||FS>60.0||SOR>4.0||
    MQRankSum<-12.5||ReadPosRankSum<-8.0" --filter-name "SNP_FILTER" -R <ref> -V <
    merged.gvcf> -O <filt.vcf>

gatk SelectVariants --select-type-to-include SNP -R <ref> -V <filt.vcf> -O <final.vcf>

gatk VariantsToTable -V <final.vcf> -F CHROM -F POS -F ID -F REF -F ALT -F QUAL -GF GT
     -O <gatk.tsv>
```

#### 1.3 freeBayes

```
freebayes --ploidy 1 --report-monomorphic -f <ref> -L <bam.list> > <fb.vcf>

vcffilter -f "QUAL>20␣&␣TYPE=snp" <fb.vcf> > <fb.filt.vcf>

get_snp_freebayes <fb.filt.vcf> > <fb.tsv>
```

#### 1.4 Samtools

```
samtools view -h <input.bam> | awk '$1␣~/^@/␣||␣($1␣!~␣/^@/␣&&␣$5>=30)' | samtools
    view -b > <sm.filt.bam>

bcftools mpileup  -O u -f <ref> -b <sm_bam.list> | bcftools call -mv -O v -o <sm.vcf>

gatk SelectVariants -V <sm.vcf> --select-type-to-include SNP -O <sm_snp.vcf>

gatk VariantsToTable -V <sm_snp.vcf> -F CHROM -F POS -F ID -F REF -F ALT -F QUAL -GF
    GT -O <samtools.tsv>
```

#### 1.5 BactSNP

```
bactsnp -q <samp.list> -r <ref> -o <output>
```

#### 1.6 VarScan

```
samtools mpileup -f <ref> -b <bam.list> -o <varscan.mpileup>

varscan mpileup2cns <varscan.mpileup> -vcf-sample-list <sample_varscan.list> > <
    varscan.out>
```

#### 1.7 Breseq

```
breseq -j 16 --brief-html-output  -r <ref> -o <tem_out> <in1.fq> <in2.fq>

bcftools view -Oz -o <tem_out>/<output.vcf.gz><tem_out>/<output_raw.vcf>

bcftools index <tem_out>/<output.vcf.gz>

bcftools merge -o <combined.vcf> -O v <tem_out_strain1>/<output_raw.vcf> <
    tem_out_strain2>/<output_raw.vcf> ...

gatk SelectVariants --select-type-to-include SNP -R <ref> -V <combined.vcf> -O <final.
    vcf>

gatk VariantsToTable -V <final.vcf> -F CHROM -F POS -F ID -F REF -F ALT -F QUAL -GF GT
     -O <breseq.tsv>
```

##### 1.8 Snippy

```
snippy --outdir 'strain1' --R1 <s1_in1.fq> --R2 <s1_in2.fq> --ref <ref> --minfrac 0.9

snippy --outdir 'strain2' --R1 <s2_in1.fq> --R2 <s2_in2.fq> --ref <ref> --minfrac 0.9

...

snippy-core --ref <ref> <strain1> <strain2> ...
```

#### 2 Supplementary Results

##### 2.1 AccuSNV shows good robustness in simulated contaminated samples

To evaluate the robustness of SNV calling tools under contamination, we generated four sets of simulated datasets using two contamination scenarios: (1) contamination from other strains of the same species, and (2) contamination from closely related species. For each scenario, we tested at two sequencing depths: 50X and 20X. Specifically, for same-species contamination, we simulated a primary isolate of *E. coli* at either 50X or 20X coverage and mixed it with a second contaminating *E. coli* isolate simulated at 10X coverage. For cross-species contamination, we used a similar approach, mixing a high-depth sample from a *Staphylococcus aureus* isolate (50X or 20X) with a 10X sample from a *Staphylococcus epidermidis* isolate as contamination. We varied the number of contaminated samples in the dataset (1, 3, 5, or all) and benchmarked the performance of each tool across these settings using precision, recall, and F1-score.

When the contamination originates from other strains of the same species, most tools suffer a substantial decrease in recall, especially under the 20X condition (Supplementary Figure S6 and S7). However, AccuSNV still preserves competitive F1-scores due to its ability to retain high precision. In the case of contamination from different species, VarScan shows high precision at 50X but experiences degraded recall and F1-score at 20X (Supplementary Figure S8 and S9). By contrast, AccuSNV maintains balanced performance across both precision and recall in all tested scenarios.

In addition, AccuSNV consistently achieves high precision and competitive F1 scores across all contamination levels. In particular, under low-depth (20X) conditions and high contamination levels, AccuSNV maintains notably higher precision and F1-scores than other tools. While tools such as GATK, Samtools, and BactSNP exhibit sharp performance drops as the level of contamination increases, AccuSNV remains comparatively robust.

These results demonstrate the effectiveness of AccuSNV in handling noisy and complex contamination patterns that commonly arise in real-world whole genome sequencing data.

##### 2.2 Ablation study on the normalization-based adjustment

Because the dataset used in “Experiments on highly variable isolates” of the main article contains a large number of isolates with different mutation rates and heterogeneous depths, it can contain extreme “gap cases”, where the alternative allele is supported by orders-of-magnitude fewer reads than the major allele (Supplementary Figure S5). To assess the contribution of AccuSNV’s normalization-based adjustment (see section “Feature extraction” in the main article) that specifically addresses such cases, we additionally evaluated a CNN-only variant without this adjustment.

For the CNN-only variant of AccuSNV, the performance matched the standard AccuSNV at the lowest mutation rate ($5 \times 10^{-10}$). However, as mutation rates increased, its precision decreased relative to the standard AccuSNV, reflecting its susceptibility to gap cases. In contrast, standard AccuSNV maintained robustness to extreme coverage imbalance, reducing false positives while preserving high recall, thereby highlighting the advantages of the current architecture.

### 3 Supplementary Figures

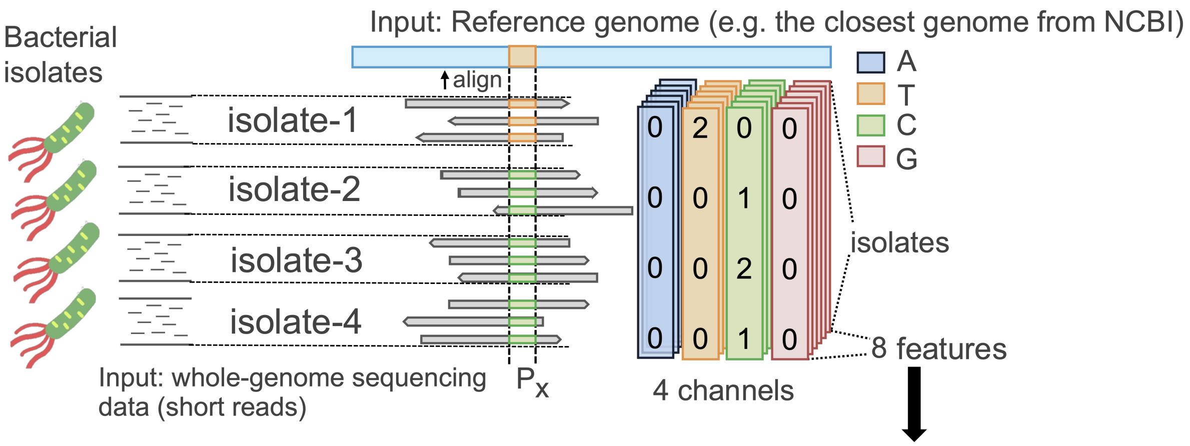


| Feature name (index in the input tensor) | Description |
|---|---|
| $d_{i,j}^{\text{norm\_fwd}}$ **(0)** | Forward-strand normalized depth for sample $i$ at locus $j$: $d_{i,j}^{\text{norm\_fwd}} = \frac{d_{i,j}^{\text{fwd}}}{\text{median}(D_i)}$, where $d_{i,j}^{\text{fwd}}$ is the forward-strand read depth and $\text{median}(D_i)$ is the median depth across all loci in sample $i$. |
| $d_{i,j}^{\text{norm\_rev}}$ **(1)** | Reverse-strand normalized depth for sample $i$ at locus $j$: $d_{i,j}^{\text{norm\_rev}} = \frac{d_{i,j}^{\text{rev}}}{\text{median}(D_i)}$, where $d_{i,j}^{\text{rev}}$ is the reverse-strand read depth; the same per-sample median depth $\text{median}(D_i)$ is used as the denominator. |
| $r_{i,j}^{\text{fwd}}$ **(2)** | Forward-strand relative depth across isolates at locus $j$: $r_{i,j}^{\text{fwd}} = \frac{d_{i,j}^{\text{fwd}}}{\sum_{x=1}^{n} d_{x,j}^{\text{fwd}}}$, capturing the share of forward-strand coverage contributed by sample $i$ among all $n$ samples. |
| $r_{i,j}^{\text{rev}}$ **(3)** | Reverse-strand relative depth across isolates at locus $j$: $r_{i,j}^{\text{rev}} = \frac{d_{i,j}^{\text{rev}}}{\sum_{x=1}^{n} d_{x,j}^{\text{rev}}}$, the reverse-strand analogue of the above. |
| **Depth_forward (4)** | Raw forward-strand depth, broadcast to the four base channels per sample. |
| **Depth_reverse (5)** | Raw reverse-strand depth, broadcast to the four base channels per sample. |
| **Qual (6)** | Base call quality score, broadcast to the four base channels per sample. |
| **Indel (7)** | The number of reads supporting an insertion and the number supporting a deletion, broadcast to the four base channels per sample. |

Supplementary Figure S1: **Description of all features used by the model**

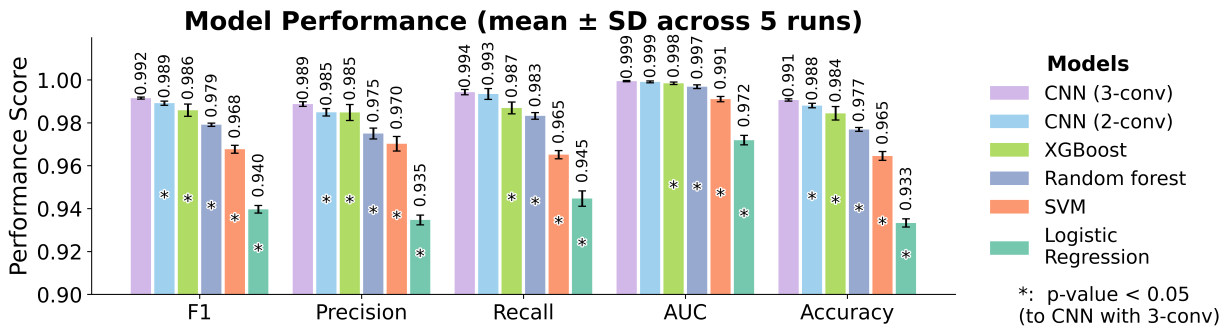


Supplementary Figure S2: **Performance comparison of CNN and other machine learning models on the real-world validation dataset.** The CNN with three convolutional layers (CNN 3-conv) achieves the highest performance across all evaluation metrics, significantly outperforming alternative machine learning models including XGBoost, random forest, SVM, and logistic regression. *: statistically significant differences (p-value < 0.05) compared to the CNN (3-conv) model using one-tailed Wilcoxon signed-rank test. Error bars represent standard deviations, demonstrating the consistency and reliability of the CNN (3-conv) approach across multiple runs.

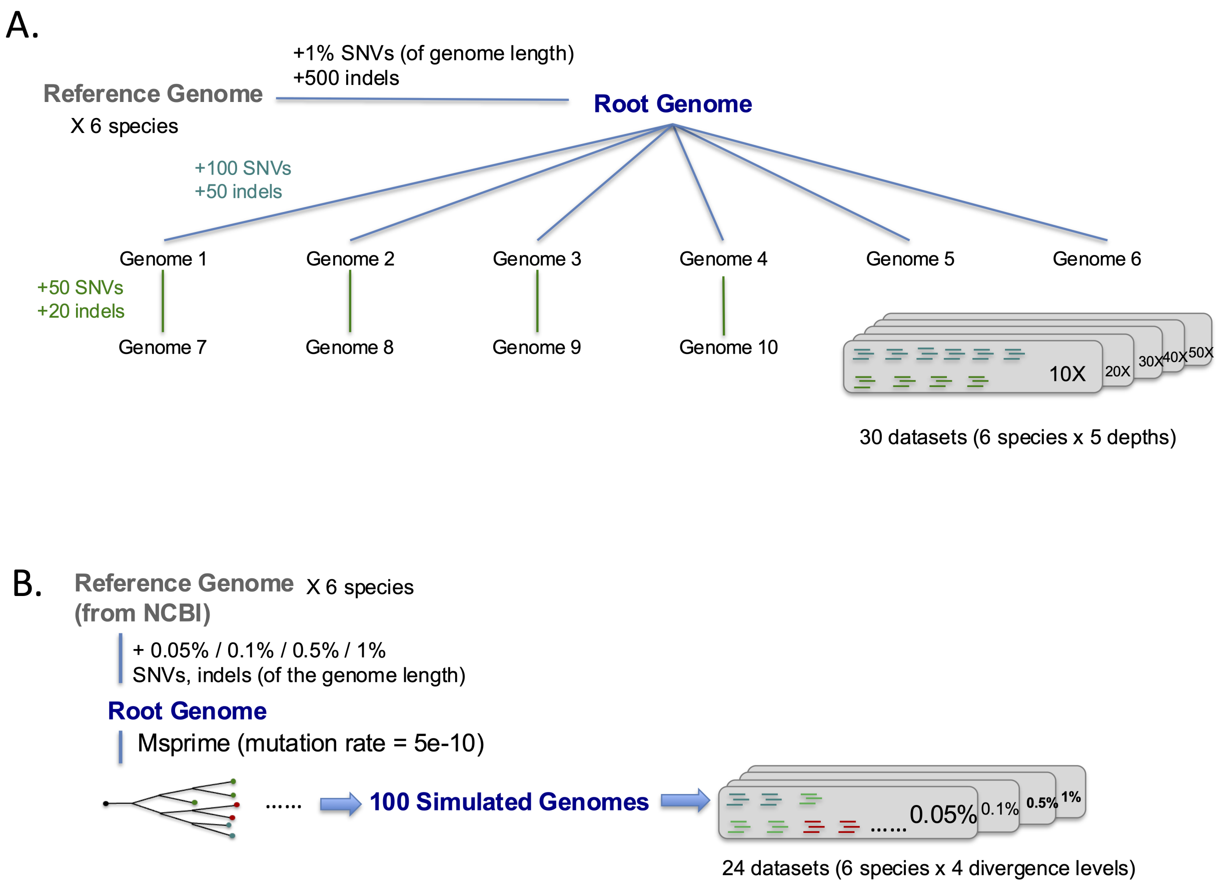


Supplementary Figure S3: **Overview of simulation designs used to benchmark SNV calling tools.** **(A)** To compare performance across sequencing depths, we simulated datasets at 10X, 20X, 30X, 40X, and 50X coverage for six representative bacterial species (*C. acnes*, *C. difficile*, *E. coli*, *K. pneumoniae*, *S. aureus*, and *S. pneumoniae*). For each species, we first introduced 1% SNVs and 500 indels into a reference genome using SimuG to generate a synthetic "root genome." Ten related genomes were then created per species by introducing additional mutations in a structured phylogeny, mimicking realistic outbreak scenarios. Illumina paired-end reads were simulated from these genomes at five different depths using ART, yielding a total of 300 datasets (6 species $\times$ 10 genomes $\times$ 5 depths). **(B)** To evaluate tool performance across varying intraspecies divergence levels, we generated four root genomes per species by introducing 0.05%, 0.1%, 0.5%, or 1% SNVs and indels into the reference genome. For each root genome, a population of 100 strains was simulated using Msprime under a neutral model with a mutation rate of $5 \times 10^{-10}$, followed by mutation assignment (80% SNVs, 20% indels) and genome generation with SimuG. Paired-end reads were simulated at random depths (15X–70X). This design produced 2,400 datasets in total (100 strains $\times$ 6 species $\times$ 4 divergence levels), enabling evaluation of SNV callers under diverse evolutionary scenarios.

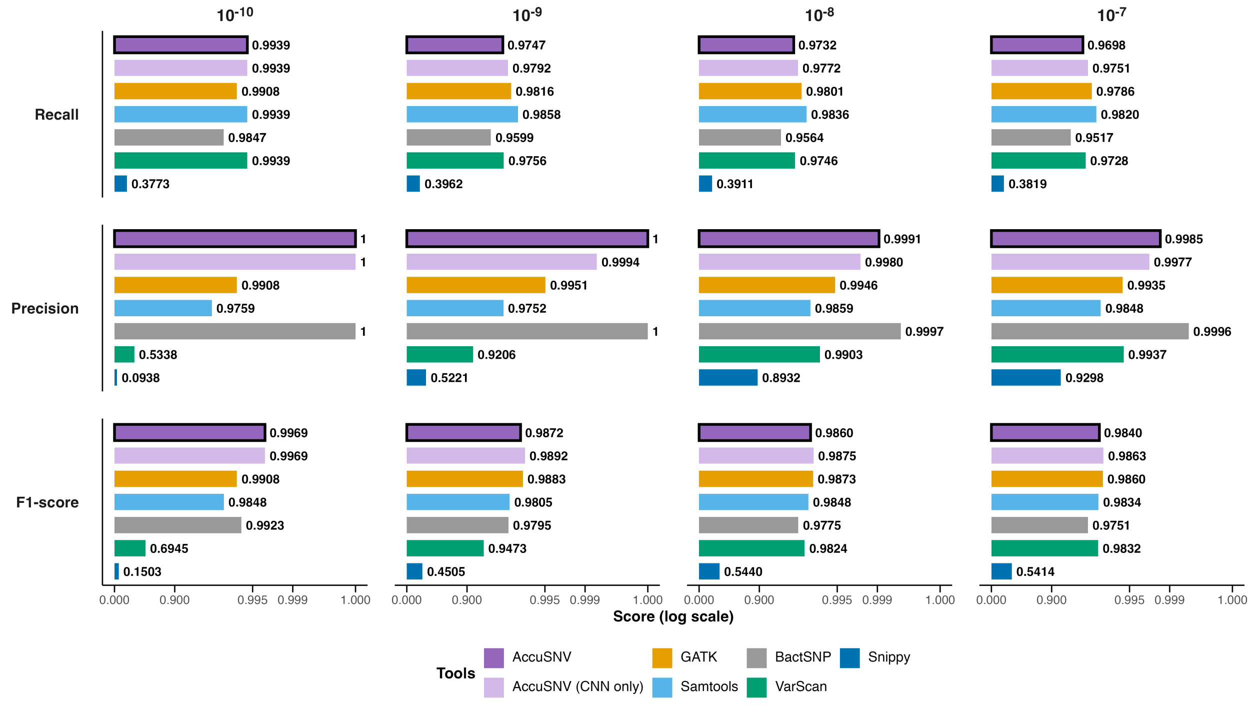


Supplementary Figure S4: **Performance comparison of SNV calling tools across datasets with increasing mutation rates between isolates.** We simulated four *E. coli* datasets with varying mutation rates ($5 \times 10^{-10}$, $5 \times 10^{-9}$, $5 \times 10^{-8}$, and $1 \times 10^{-7}$, mutations per base per generation), labeled as "$10^{-10}$", "$10^{-9}$", "$10^{-8}$", and "$10^{-7}$", respectively, to evaluate the robustness of different SNV calling tools on genetically diverse bacterial isolates. Panels show average recall (top), precision (middle), and F1 score (bottom). AccuSNV bars are highlighted with a black outline. AccuSNV (purple) denotes the standard version with the normalization-based adjustment integrated into feature encoding, while "AccuSNV (CNN only)" (light purple) represents the ablation variant without this adjustment. This ablation study is shown only for the highly variable isolates dataset, where heterogeneous depth and frequent gap cases provide the most stringent conditions to evaluate the contribution of the adjustment.

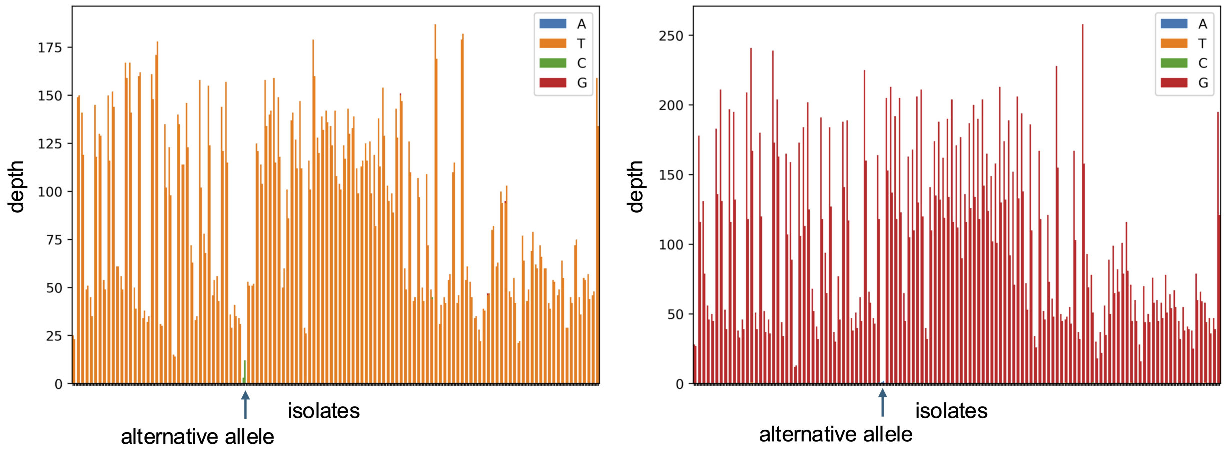


Supplementary Figure S5: **Examples of gap cases caused by extreme depth imbalance.** Depth distributions of nucleotides across isolates are shown for two candidate SNV positions. Each bar represents the read depth of a specific nucleotide (A, T, C, or G) in one isolate. In both examples, the putative alternative allele (indicated by the arrow) is supported by only a few reads, while the major allele exhibits much higher depth across nearly all isolates. Such extreme coverage imbalances ("gap cases") are rare in the training data and often lead to misclassification by AccuSNV without any adjustment.

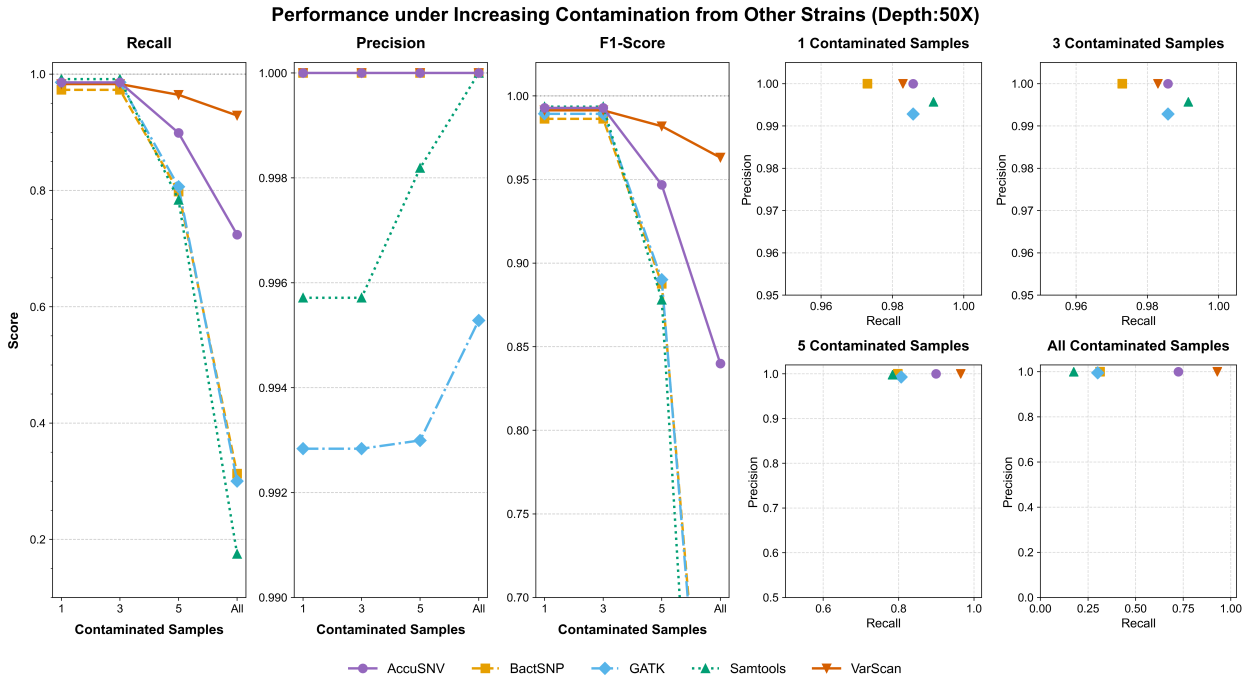


Supplementary Figure S6: **Performance under increasing contamination from other strains (depth = 50X).** This figure shows the recall, precision, and F1-score of five SNV calling tools (AccuSNV, BactSNP, GATK, Samtools, and VarScan) as the number of contaminated samples increases from 1 to all samples. Contamination was simulated by mixing one isolate's 50X clean reads with 10X reads from a different isolate of the same species. AccuSNV maintains high performance across all metrics and levels of contamination, while other tools show notable declines in recall and F1-score, especially when more samples are contaminated.

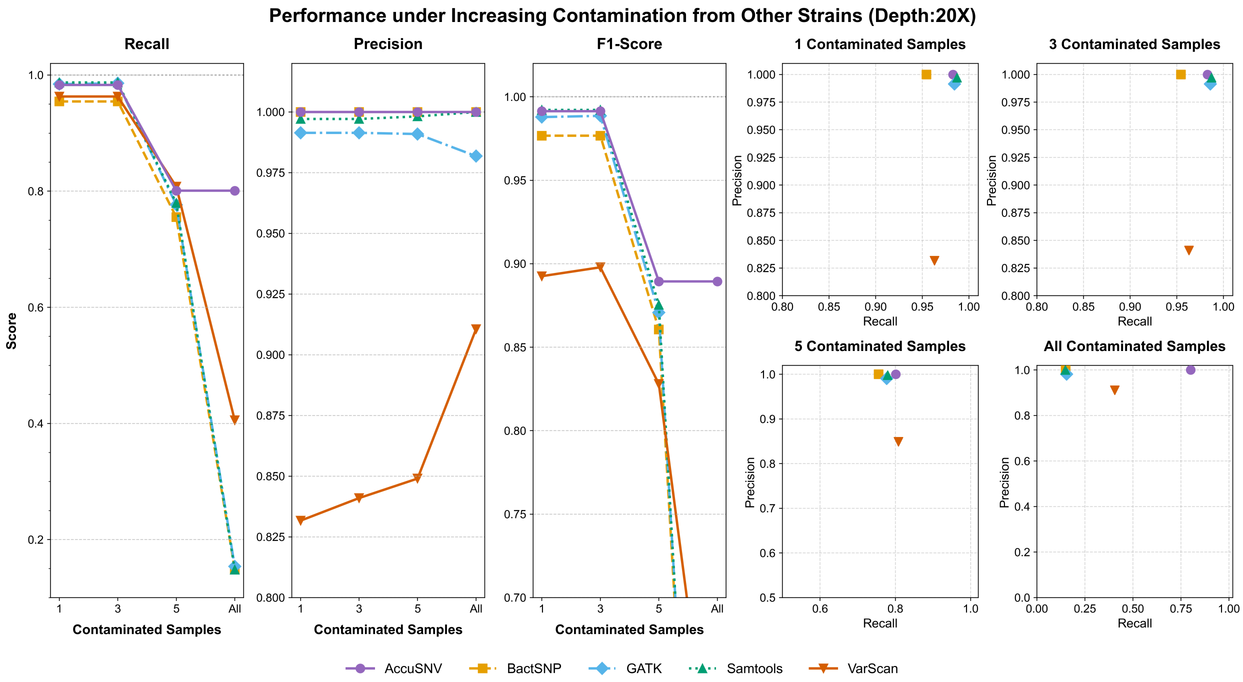


Supplementary Figure S7: **Performance under increasing contamination from other strains (depth = 20X).** Same setting as Supplementary Figure S2, except clean reads were simulated at 20X coverage to evaluate tool robustness under lower sequencing depth. Most tools, especially GATK and Samtools, experience sharp drops in recall and F1-score as contamination increases. In contrast, AccuSNV maintains stable precision and outperforms others in F1-score across contamination levels.

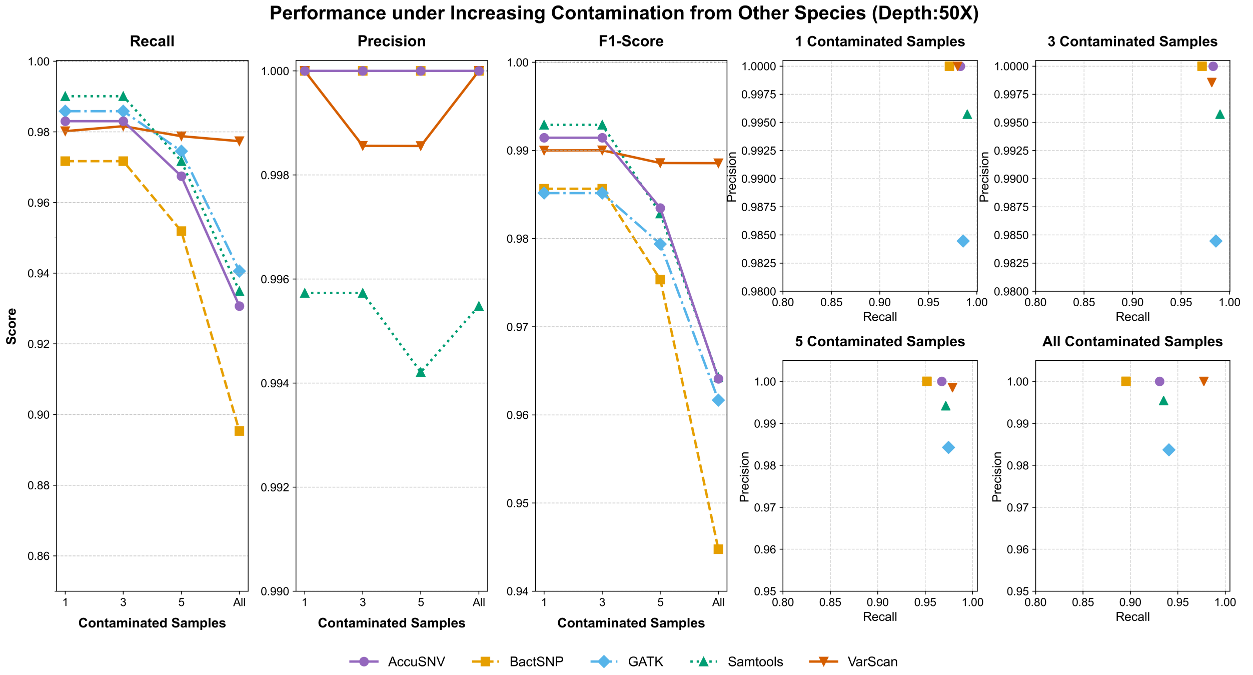


Supplementary Figure S8: **Performance under increasing contamination from other closely related species (depth = 50X).** Contamination was simulated by mixing 50X reads from one isolate of *S. aureus* with 10X reads from an isolate of *S. epidermidis*. Despite the increased complexity introduced by inter-species contamination, AccuSNV retains high precision and balanced F1-score across all contamination levels. Other tools, particularly BactSNP, exhibit reduced recall, highlighting their reduced robustness to this type of contamination.

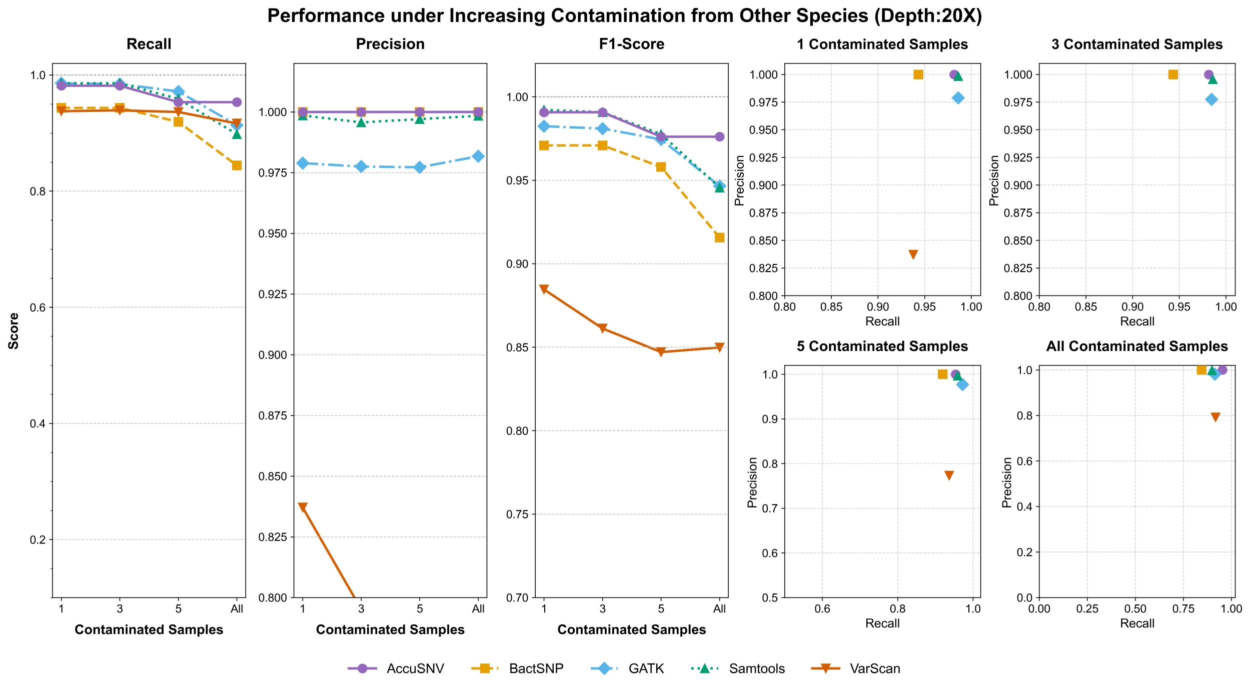


Supplementary Figure S9: **Performance under increasing contamination from other closely related species (depth = 20X).** Same setting as Supplementary Figure S4 but under 20X sequencing depth. The performance of most tools further degrades, with recall and F1-score dropping substantially as contamination increases. AccuSNV remains the most robust, achieving the highest F1-score across all contamination levels, due to its ability to balance precision and recall even in noisy and low-coverage settings.

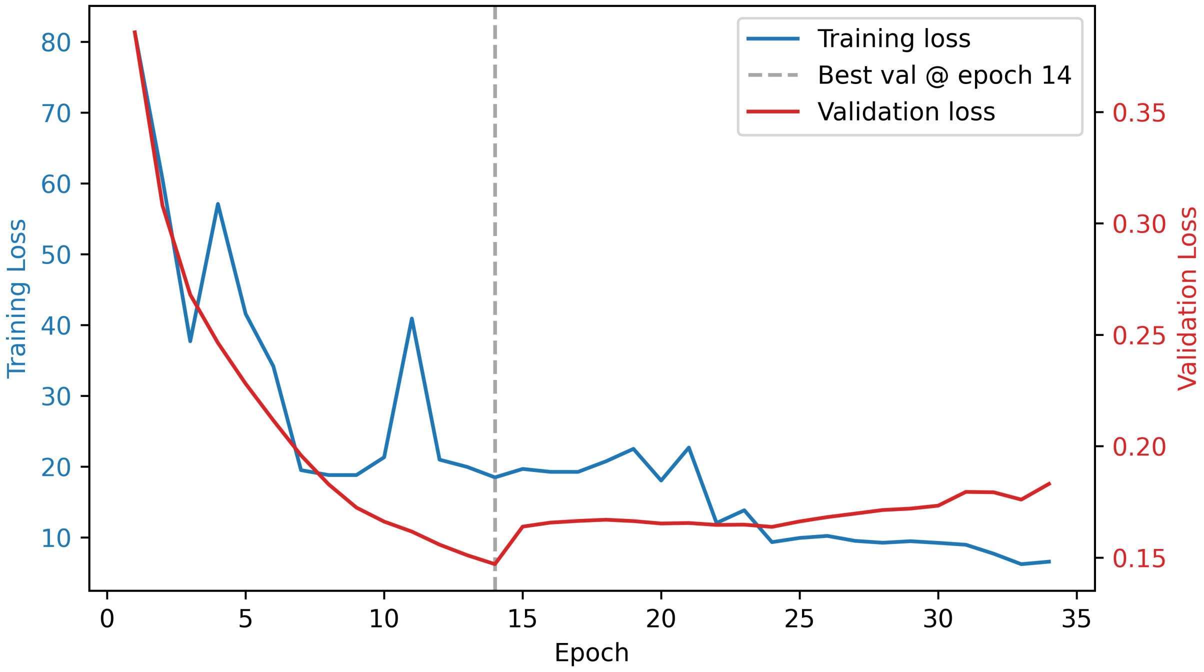


Supplementary Figure S10: **Learning curves during model training.** The training loss (blue) and validation loss (red) both decrease during early training, with the validation loss reaching its minimum at epoch 14 (gray dashed line). Subsequent increases in validation loss while training loss continues to decrease suggest the onset of overfitting beyond this point.

### 4 Supplementary Tables

| Tools | *E. coli* (n=10) | | *E. coli* (n=100) | |
|---|---|---|---|---|
| | Runtime (min) | Memory (MB) | Runtime (min) | Memory (MB) |
| AccuSNV | 30.3 | 1,140 | 64.5 | 1,340 |
| BactSNP | 13.1 | 1,722 | 340.6 | 2,017 |
| GATK | 47.6 | 1,275 | 435.5 | 1,320 |
| Snippy | 5.4 | 719 | 50.1 | 1,105 |
| Samtools | 51.3 | 1,205 | 518.4 | 1,259 |
| VarScan | 60.2 | 1,310 | 593.8 | 1,410 |
| Breseq | 464.5 | 1,099 | >2,880 | - |
| freeBayes | 922.1 | 1,443 | >2,880 | - |

Supplementary Table S1: **Computational performance comparison of SNV calling tools.** Runtime (minutes, from raw reads processing to final results) and peak memory usage (MB) for eight SNV calling tools evaluated on simulated *E. coli* datasets with 10 isolates (50X depth) and 100 isolates (15X-70X depths). Each tool was run on an AMD EPYC 7513 2.6GHz 64-core processor with 24Gb of allocated memory with 8 cores and a 48-hour timeout limit. Values marked with ">" indicate jobs that exceeded the timeout period and were terminated. "-": missing values due to terminated jobs.

| Study | Species | Description about the criteria used in each study to define “quality-filtered” SNVs |
|---|---|---|
| Zhao, 2019 | *B. fragilis* | In particular, genomic positions were considered to be candidate SNV positions if at least one pair of isolates was discordant on the called base and both members of the pair had: FQ scores (produced by SAMtools) less than -60, at least 7 reads that aligned to each of the forward strand and reverse strand, and a major allele frequency of at least 90%. If the median coverage across samples at the candidate position was less than 10 reads or if 33% or more of the isolates failed to meet the filters described above, this position was discarded. For each SNV position identified, a nucleotide call was assigned to each isolate using the major allele call across reads for that isolate at that position. If fewer than 7 reads aligned to either the forward or reverse strand of a position in an isolate, or the major allele frequency was smaller than 90%, an ambiguous call was assigned to the isolate at that SNV position. |
| Conwill, 2022 | *C. acnes* | To determine SNV positions within each lineage, basecalling was repeated using the following process:<br>first, basecalls were marked as ambiguous if the FQ score produced by SAMtools was above -30, the coverage per strand was below 3, the major allele frequency was below 0.75, or more than 25% of reads supported indels; second, genomic positions with a median coverage below 12 reads across samples or where at least 34% of basecalls were ambiguous across samples were omitted. In addition, to remove variants that emerged from recombination or other complex events, we identified SNVs that were less then 500 bases apart and for which the correlation of non-ancestral allele frequencies (see below) across colonies within a lineage exceeded 0.75 (Table S8); these positions, as well as regions on the reference genome with homology to plasmids (see C. acnes plasmid analysis), were removed from down- stream analysis.<br>All remaining genomic positions that passed these strict filters and retained two non-ambiguous alleles were considered SNV positions and were investigated across samples. To call genotypes for as many colonies as possible at these SNV positions, including ones with low coverage, basecalls were repopulated from the raw data and only marked as ambiguous only if the coverage per strand was below 1, the total coverage below 3, the major allele frequency below 0.67, or more than 25% of reads supported a deletion. Details on SNVs detected in each lineage are available in Table S6. |
| Baker, 2024 | *C. acnes* & *S. epidermidis* | To initially filter low-quality calls in the alignments to lineage co-assemblies, we considered calls with an FQ score <30, major allele frequency <.75, or coverage <4 as “N”. Next, we excluded positions where >40% of samples have an N in that position. To build phylogenies, nucleotide calls at accepted positions were less strictly filtered to retain information at these highly informative positions. Major-allele nucleotides were used as calls provided they had a major allele frequency of 70% and a coverage of at least 5 reads (forward or reverse). |

Supplementary Table S2: Summary of criteria used in published studies to obtain “quality-filtered” SNVs.
